# AURKA inhibition amplifies DNA replication stress to foster WEE1 kinase dependency and synergistic antitumor effects with WEE1 inhibition in cancers

**DOI:** 10.1101/2025.05.28.656693

**Authors:** Jong Woo Lee, Jackie Shi, Julian Barrantes, Pratima Chaurasia, Sundong Kim, Sebastian Cruz-Gomez, Cindy Yang, Jianlei Gu, Hongyu Zhao, Dejian Zhao, Benjamin Schiff, Ansley Roche, Elizabeth B. Perry, Jeffrey P. Townsend, Gary M. Kupfer, Katerina Politi, Erica Golemis, Barbara Burtness

## Abstract

Highly elevated expression of the oncogene Aurora kinase A (AURKA) occurs in numerous human cancers harboring defective p53, nominating AURKA as a potential vulnerability in *TP53*-mutated cancer. However, clinical trials have indicated modest monotherapy activity of AURKA inhibitors. Here, we demonstrate that AURKA inhibition promotes phosphorylation of Replication Protein A (RPA), resulting in stalled DNA replication fork progression and eliciting a replication stress response in multiple *TP53*-mutated models, creating a druggable dependence on the mitotic checkpoint kinase WEE1. Combined inhibition of AURKA and WEE1 synergistically enhanced replication stress, tumor-specific apoptotic cell death, and mitotic catastrophe, and lead to marked tumor regression in cell line- and patient-derived xenograft models of *TP53*-mutated cancer. Our findings define enhanced DNA replication stress as underlying the strong synergy between AURKA and WEE1 inhibitors and offer preclinical confirmation of efficacy, indicating high potential for clinical translation of this synthetic lethal strategy for *TP53*-mutated carcinomas.

**Statement of significance:** We demonstrate that a small molecule AURKA inhibitor amplifies DNA replication stress in *TP53*-mutated carcinomas. This amplification of DNA replication stress can be leveraged this for synthetic lethal therapy in a combination with WEE1 inhibition that enhances antitumor effects in *in vitro*, in xenografts and in patient-derived xenograft models, advancing a promising novel combination therapy.

## INTRODUCTION

Loss-of-function *TP53* mutations are the most common genomic alterations in human cancer and are associated with increased aggressiveness of malignancy, faster treatment resistance, and worse prognosis. Therapeutic strategies for *TP53*-mutated cancers that exploit survival pathways required in the absence of p53 function are of great interest for cancer therapy (1). We therefore sought targetable kinases regulating both DNA replication fork and mitotic entry/progression to design such synthetic lethal treatments. Initial screens identified Aurora kinase A (AURKA/STK15) and WEE1 as potential targets (2–4). Each of these kinases is known to be upregulated in the setting of *TP53* mutation, and each has a distinct role in protecting DNA replication and regulating mitosis (5–8); for each, clinical development of inhibitors is ongoing (9,10).

Aurora kinases are a family of serine/threonine kinases that are crucial to mitotic entry, cell division and cell survival in all eukaryotes. As noted, within this family, AURKA and p53 reciprocally regulate each other through a number of interactions: among these, p53 directly inhibits AURKA kinase activity (11), while p53 phosphorylation by AURKA at serine 315 facilitates p53 ubiquitination and degradation (12), and p53 phosphorylation at serine 215 inhibits p53 DNA-binding and transactivation (13). AURKA plays pleiotropic roles in: (a) centrosome maturation through activating phosphorylation of the polo-like kinase 1 (PLK1; ref. 14), (b) fostering mitotic entry from G_2_/M transition by phosphorylation of the phosphatase cell division cycle 25B/C (CDC25B/C) resulting in dephosphorylation of cyclin-dependent kinase 1 (CDK1) to activate the CDK1/cyclin B (CCNB) complex (15,16), and (c) spindle assembly through interaction with TPX2 microtubule nucleation factor (17,18). Recently, evidence has emerged that AURKA functions throughout the cell cycle. For example, AURKA activation in G_1_ controls cilium disassembly (19) and the TPX2/AURKA heterodimer binds to the DNA repair protein 53BP1, stabilizing replication forks (7). Despite these and other critical roles of AURKA in cell division and survival, clinical activity of single-agent AURKA inhibition has been modest, with response rates of <10% observed for the first-generation inhibitor alisertib (MLN8237) in solid tumors (20,21).

Specific mechanisms of resistance to AURKA inhibition have been previously demonstrated (22). In *TP53*-mutated head and neck squamous cell carcinomas (HNSCCs), we found that AURKA inhibition results in cell cycle arrest mediated by WEE1, long known as a G_2_/M checkpoint kinase but more recently recognized for a key role in replication stress response (23,24). WEE1 restrains mitotic entry through an inhibitory phosphorylation of CDK1 at tyrosine 15 and phosphorylates CDK2 to delay G_1_/S transition; thus, its inhibition interferes with cell cycle checkpoints that are central to the replication stress response (8,24,25).

Replication stress-the disruption of DNA replication progression and stalling of replication forks-is an important cause of genomic instability in cancer and constitutes a potential therapeutic vulnerability to cancer treatment (26). AURKA inhibition reduces replication fork speed and increases transcription-replication conflicts (7); these changes activate the ataxia telangiectasia and Rad3-related (ATR) kinase, ATR activates checkpoint kinase 1 (CHK1), and CHK1 in turn phosphorylates WEE1 (27). WEE1 activation stabilizes replication forks through effects on checkpoint kinase 2 (CHK2) and the 53BP1 pathway. Together, these activities limit replication stress-induced DNA double strand breaks and result in S phase and G_2_/M boundary cell cycle and mitotic delay (8,28).

WEE1 inhibitors have been proposed for cancers with high endogenous replication stress, as reflected in elevated Cyclin E (CCNE1; refs. 29-31). Combinations of WEE1 inhibition with replication stress-inducing chemotherapy agents have been proposed, but these combinations may be limited by overlapping toxicity (32,33), again highlighting the need for novel, optimized mechanism-based combination approaches. Cell cycle arrest apprears to be the intrinsic limitation of AURKA inhibition monotherapy. Combination of the WEE1 inhibitor adavosertib with alisertib causes dephosphorylation of the cell-cycle inhibitory phosphorylation of CDK1. This disruption of cell cycle arrest results in mitotic catastrophe and cell death, producing synergistic antitumor effects in *in vitro* and *in vivo* HNSCC models (22). However, the manner in which AURKA induced WEE1 to produce the conditions for synergistic inhibition of these two targets has not been understood. Here, we reveal the role of replication stress in fostering this synergy in the *p53*-mutated context, and provide evidence that this effect is general across multiple *TP53*-mutated solid cancers.

Given the complementary respective roles of AURKA and WEE1 in stabilizing the DNA replication fork and responding to replication stress in S phase to coordinate mitotic delay, we explored the interplay of these effects in producing synergistic anti-cancer effects, demonstrating that AURKA inhibition produces a dramatic increase in replication stress, thereby inducing a replication stress response program that activates WEE1, thus creating WEE1 dependency and the conditions for synergy through dual AURKA and WEE1 inhibition. Employing the novel and highly AURKA-specific inhibitor VIC-1911 (previously TAS-119; Vitrac Therapeutics), we show synergy for the combination of VIC-1911 and WEE1 inhibition across a series of *TP53*-mutated human cancer cell lines including HNSCC, lung, pancreatic, ovarian and hepatocellular cancer, as well as xenograft and patient-derived xenograft (PDX) HNSCC and lung cancer models. Our findings indicate high therapeutic potential for dual inhibition of AURKA and WEE1.

## RESULTS

### Concomitant VIC-1911 inhibition of AURKA with advavosertib inhibition of WEE1 synergistically inhibits cell growth and survival in human cancers

As AURKA is highly overexpressed in *TP53*-mutated human cancers and its expression is associated with poor outcome. Accordingly, we initially screened protein expression of AURKA and its coactivator TPX2 in a panel of *TP53*-mutated HNSCC (FaDu, CAL27, Detroit 562, UNC7, SCC9 and SCC61) and lung cancer (NCI-H520, NCI-H1437, NCI-H358, NCI-H1792, NCI-H1734 and A549) human cancer cell lines. AURKA protein content was dramatically increased in all tested cancer cells relative to both normal human tracheobronchial epithelial cells (NHTBE) and normal human fibroblasts (BJ1). Further, as expected, expression of the AURKA partner/activator TPX2 strongly correlated with AURKA expression level (Supplementary Fig. S1A). As well, we confirmed that *AURKA* expression is elevated in lung cancers including squamous cell lung carcinoma (SCLC) and lung adenocarcinoma (LUAD) as determined from Oncomine (three individual gene expression datasets; *P* < 0.0001) and its overexpression is associated with worse overall survival in lung cancer patients using Kaplan-Meier plotter analysis with a platform based on the GEO, EGA and TCGA databases (logrank test *P* =3 × 10^-4^ in LUAD; *P* = 0.033 in SCLC; Supplementary Fig.S1B and S1C; ref. 34).

The AURKA inhibitor alisertib and WEE1 inhibitor adavosertib are synergistic against HNSCC tumors (22). To explore whether this synergy IS disease-specific or present in a range of *TP53*-mutated solid tumors, we tested drug sensitivities to either adavosertib or VIC-1911 in ovarian carcinoma (OVC), pancreatic ductal adenocarcinoma (PDAC), and hepatocellular carcinoma cells (HCC). With the HNSCC and lung cancer lines above, we have data in a total of 17 cells lines, of which FaDu, CAL27, Detroit 562, UNC7, SCC9, SCC61, NCI-H520, NCI-H358, NCI-H1437, NCI-H441, NCI-H1792, NCI-H23, SK-OV-3 and MiaPaCa-2 cell lines are *TP53*-mutated, and A549 and A2780 cell lines are *TP53* WT. Most cancer cells were sensitive to each drug given alone, with IC_50_ of 162 to 949 nmol/L, whereas some cancer cells were relatively resistant to one or both agents. For adavosertib, these were HNSCC SCC61 (IC_50_ = 1.26 µmol/L) and ovarian cancer A2780 (IC_50_ = 1.12 µmol/L) cell lines (Fig. 1A), and for VIC-1911, HNSCC UNC7 (IC_50_ = 2.13 µmol/L) and ovarian cancer SKOV-3 (IC_50_ > 2.5 µmol/L) cell lines were resistant (Fig. 1B). To see combination effects of adavosertib and VIC-1911 in a variety of cancers, we applied 8 × 8-point-dose combinations for cell viability assays in these 18 cancer cell lines treated with control, adavosertib, VIC-1911 or combination over four days. Synergy of the combination was determined by multiple validation methods, including cooperative correlation, Loewe synergy score (LSS) and dose-response curves (Fig. 1C, 1D and Supplementary Fig. S2-S4).

**Figure 1.**
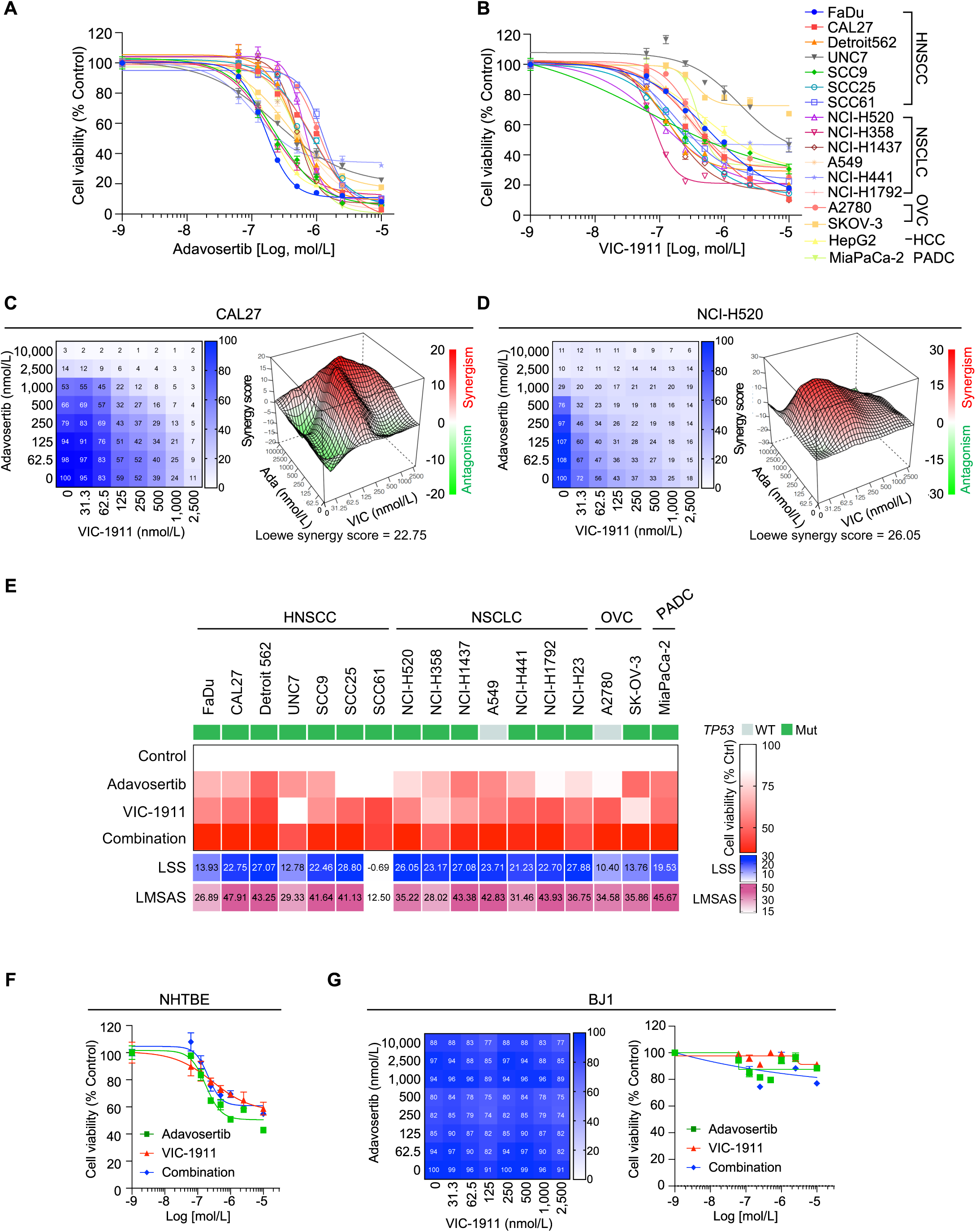
Combination drug testing reveals synergistic suppressive effect on cell viability by combined a highly selective AURKA inhibitor, VIC-1911 in combination with WEE1 inhibition in human cancer cells. **A** and **B**, Dose-response curves for determining the sensitivity of various human cancer cell lines to either adavosertib (**A**) or VIC-1911 (**B**) with multiple dose range (adavosertib: 0.0625-10 μmol/L; VIC-1911: 0.0313-2.5 μmol/L). Cell viability was determined after 4-day exposure to drugs by CellTiter-Glo assay. Cell viability was normalized to control cells. Data are shown as mean ± SEM. **C and D**, Representative images of cooperative correlation matrix (right) and Loewe plots (left) of HNSCC CAL27 cells (**C**) and NSCLC NCI-H520 cells (**D**) in combinational treatment VIC-1911/adavosertib. **E**, Status of *TP53* mutation, Loewe synergy scores (LSS) and maximum synergistic area scores (LMSAS) of human cancer cell lines are shown for evaluating synergy of combinational VIC-1911 and adavosertib treatment using SynergyFinder. and synergy score >10 and red area indicate synergism. **F** and **G**, Dose-response curves of NHTBE (**F**) and BJ1 (**G**) cells treated with control, adavosertib, VIC-1911 or combination from 4-day CellTiter-Glo assay. Cell viability was normalized to control cells. Shown are the means of technical triplicates from one experiment and the data as shown mean ± SEM are representative of three independent experiments with consistent results.

Notably, 16 of the 17 cancer cell lines showed synergistic suppressive effect of adavosertib/VIC-1911 combination on cell viability, as shown by LSS and Loewe maximum synergy area scores (LMSAS), with scores of 10.4 - 27.9 and 26.9 - 47.9, respectively. (Fig. 1E). We found that non-TP53 mutant cancer cells also exhibited some degree of synergy in responsiveness to combination treatment with LSS of 10.40 for A2780 cells or 23.71 for A549 cells, and both cell lines harbor TP53 family gene, *TP63* mutations. Cancer cells resistant to either of the single agents or to both drugs nonetheless showed synergism in most models, although only an additive effect (synergy score: -0.69) without synergy was observed in HNSCC SCC61 cells (Fig. 1E and Supplementary Fig. S2F). Additionally, synergy was also confirmed by Chou-Talalay-conjugated isobologram analysis for HNSCC cells, including FaDu, Detroit 562, CAL27 and UNC7 (CI < 1 and at least 75% synergy (effective level; Fa 0.75; Supplementary Fig. S5)). Importantly, we evaluated the off-target cytotoxicity of the combination, and neither single agent or combination treatment was cytotoxic toward either normal NHTBE or BJ1 cells (Fig. 1F and 1G).

Upregulation of AURKA is associated with poor prognosis in HPV-negative HNSCC and in lung cancer. Accordingly, we focused on these two difficult malignancies for further study (Supplementary Fig. S1C, ref. 22). Using maximum synergistic concentrations of adavosertib and VIC-1911, we assessed the inhibitory effect of the combination on clonogenic survival, and on anchorage-independent cell growth in a 3-dimensional (3D) setting. Combined administration of adavosertib and VIC-1911 inhibited clonogenic survival by more than 80% in multiple cancer cell lines (Fig. 2A). Furthermore, the combination treatment was more effective than was control or either single agent, yielding significantly fewer and smaller colonies in both soft agar and oncosphere formation assays (Fig. 2B-2D).

**Figure 2.**
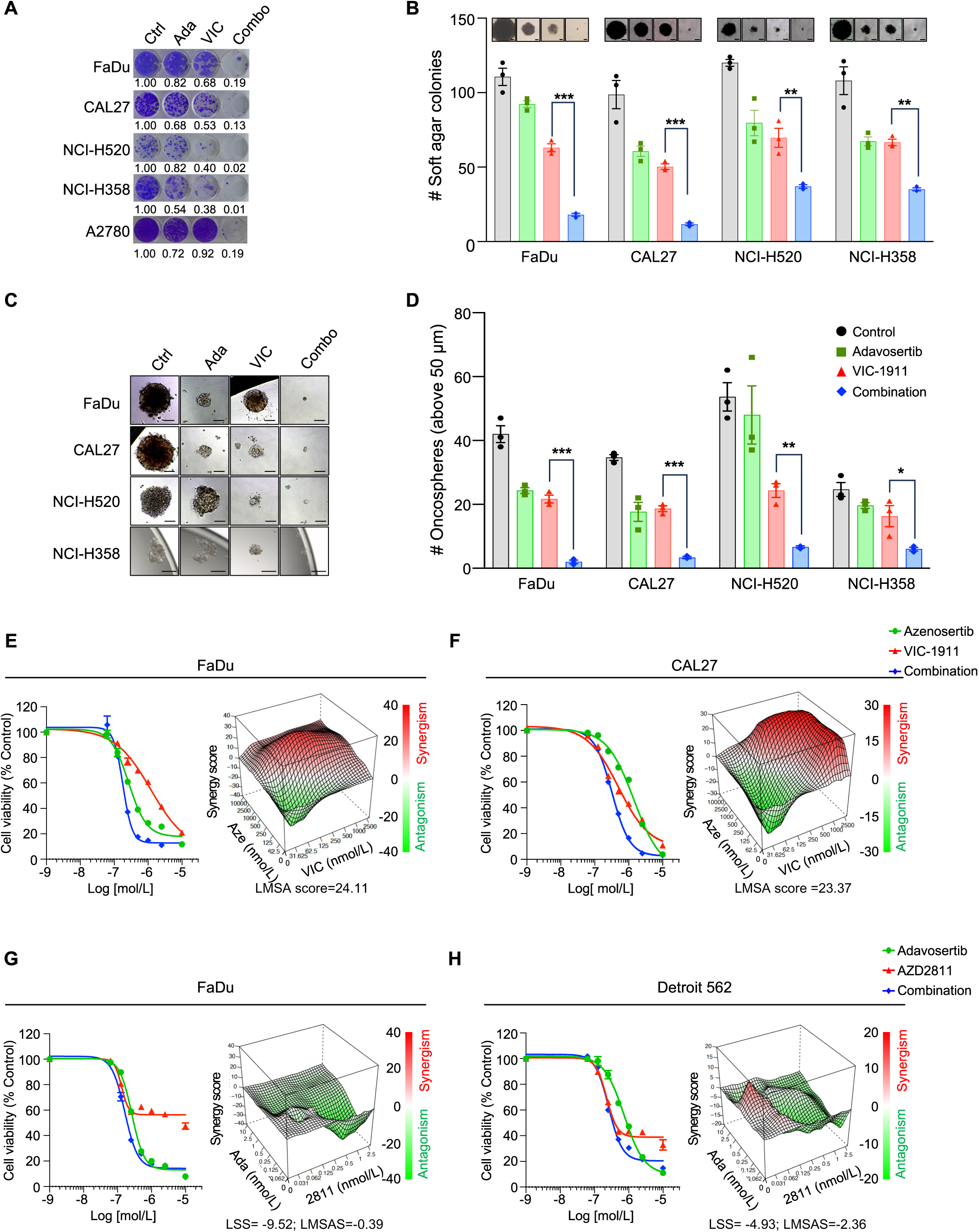
Combined AURKA/WEE1 inhibition cooperatively suppresses cell survival in HNSCC and lung cancer cells, with less cytotoxicity in human normal epithelial and fibroblast cells. **A**, Suppressive effect on clonogenic survival of VIC-1911/adavosertib combination in HNSCC (FaDu and CAL27), NSCLC (NCI-H520 and NCI-H358) and ovarian cancer (A2780) cells. Cells were treated with control, adavosertib (all other cell lines 250 nmol/L; CAL27 500 nmol/L), VIC-1911 (all other cell lines 125 nmol/L; CAL27 250 nmol/L) or combination for 4 days and cultured without drugs without drugs up to 9-12 days. Clonogenic survival was determined via crystal violet staining. Quantification of cell density from crystal violet staining is shown after normalized to control. **B**, Anchorage independent cell survival of HNSCC and lung cancer cells in response to VIC-1911/adavosertib combination. Cells were treated with the indicated drugs in low density of soft agarose cell culture setting for 4 days and cultured without drugs up to 25-30 days. A colony was defined as containing more than 10 cells, as indicated by >50 pixels in image J. Representative images are viable colony in soft agar. Scale bar = 100 μm. **C** and **D**, FaDu, CAL27, NCI-H520 and NCI-H358 cells were cultured with the conditioned media for oncosphere formation assay after 7 to 9 days of 4-day treatment with the indicated drugs as single or combination. Representative image of oncospheres (**C**) and quantification of number of oncospheres (**D**) are shown. Scale bar = 100 μm. Each dot shows a biological replicate (three/condition) and data are shown as mean ± SEM. Statistical significance was determined by Student *t* test. \**P* < 0.05; \*\**P* <0.01; \*\*\**P* < 0.001. **E** and **F**, Synergistic antitumor effect of late generation of WEE1 inhibitor azenosertib and VIC-1911 combination in HNSCC FaDu (**E**) and CAL27 (**F**) cells. Azenosertib doses are the same as adaovsertib. Shown are the means of technical triplicates from one experiment and the data as shown mean ± SEM are representative of triplicated and three independent experiments with consistent results.

Taken together, combined inhibition of AURKA and WEE1 by the highly selective AURKA inhibitor VIC-1911 and adavosertib synergistically suppressed cell growth and survival in both 2D and 3D cell culture conditions in a variety of human cancer cell lines. We further confirmed the value of dual AURKA and WEE1 inhibition in HNSCC cells using a chemically distinct, more selective WEE1 inhibitor azenosertib (Zn-c3; ref. 10), currently in clinical testing. To test for synergy, we conducted cell viability assays after treatment with control, VIC-1911, azenosertib or combination for four days followed by determination of LSS and dose-response curves in FaDu and CAL27 HNSCC cells. Similar to our findings with adavosertib, azenosertib alone showed some degree of suppressive effect on cell viability in FaDu and CAL27 cells but azenosertib combined with VIC-1911 resulted in a more pronounced suppressive effect, mirroring observations with the combination of adavosertib and VIC-1911, with LMSAS of 24.1 and 23.4 in FaDu and CAL27 cells, respectively (Fig. 2E and 2F).

### AURKA inhibition enhances DNA replication stress to induce WEE1 dependency

Having confirmed that effects are achieved across multiple p53-deficient cancer models with VIC-1911, we sought to determine the mechanism by which combination AURKA/WEE1 inhibition produces this synergy. Because of the role WEE1 plays in responding to replication stress, we explored how AURKA inhibition impacts replication fork integrity and progression. As noted, the AURKA/TPX2 complex, by negatively regulating 53BP1, plays an important role in protecting replication fork stability in S phase (7), while WEE1 (together with CHK1) is a critical replication stress response protein, suppressing CDK2 activity (7,8,25). We therefore assessed the effect of inhibiting AURKA alone or in combination with WEE1 on replication fork progression. We employed DNA fiber assays using chase-labelling with 5-chloro-2-deoxyuridine (CIdU) and 5-iodo-2-deoxyuridine (IdU) following treatment with control, VIC-1911, adavosertib or combination for 24h (Fig. 3A). AURKA and WEE1 inhibition were each individually effective in reducing replication fork speed in HNSCC and lung cancer cells, as reflected by shorter IdU-labelled DNA fibers relative to control cells in the DNA fiber assay, with AURKA inhibition having the greater effect in FaDu cells (adavosertib vs VIC-1911; *P* = 0.0002). Effects were similar in NCI-H358, CAL27, and NCI-H520 cells (Fig. 3B, 3C and Supplementary Fig. S6A-S6D). Furthermore, concomitant treatment with VIC-1911 and adavosertib further significantly delayed DNA fork progression in FaDu and NCI-H358 cells compared to VIC-1911 (FaDu, *P* = 0.0014; H358, *P* < 0.0001) or adavosertib (FaDu and NCI-H358, *P* < 0.0001) as a single agent (Fig. 3B and 3C). Notably, no effects of single or combination treatment of VIC-1911 or adavosertib on replication progress were observed in SCC61 cells, the single cell line in which we failed to confirm synergy for combined AURKA and WEE1 inhibition in our initial cell viability screening (Supplementary Fig. S6C), further supporting the central role of replication stress induction in creating synergy for this combination. To understand the lesser activation of replication stress in SCC61 cells, we performed whole-exome sequencing and identified *TP53* (p.R110L), and gene mutations of interest potentially related to AURKA and WEE1 activities, including *AJUBA* (p.E310Q) and *CDC25C* (p.G224R; p.G254R; p.G263R; p.G297R).

**Figure 3.**
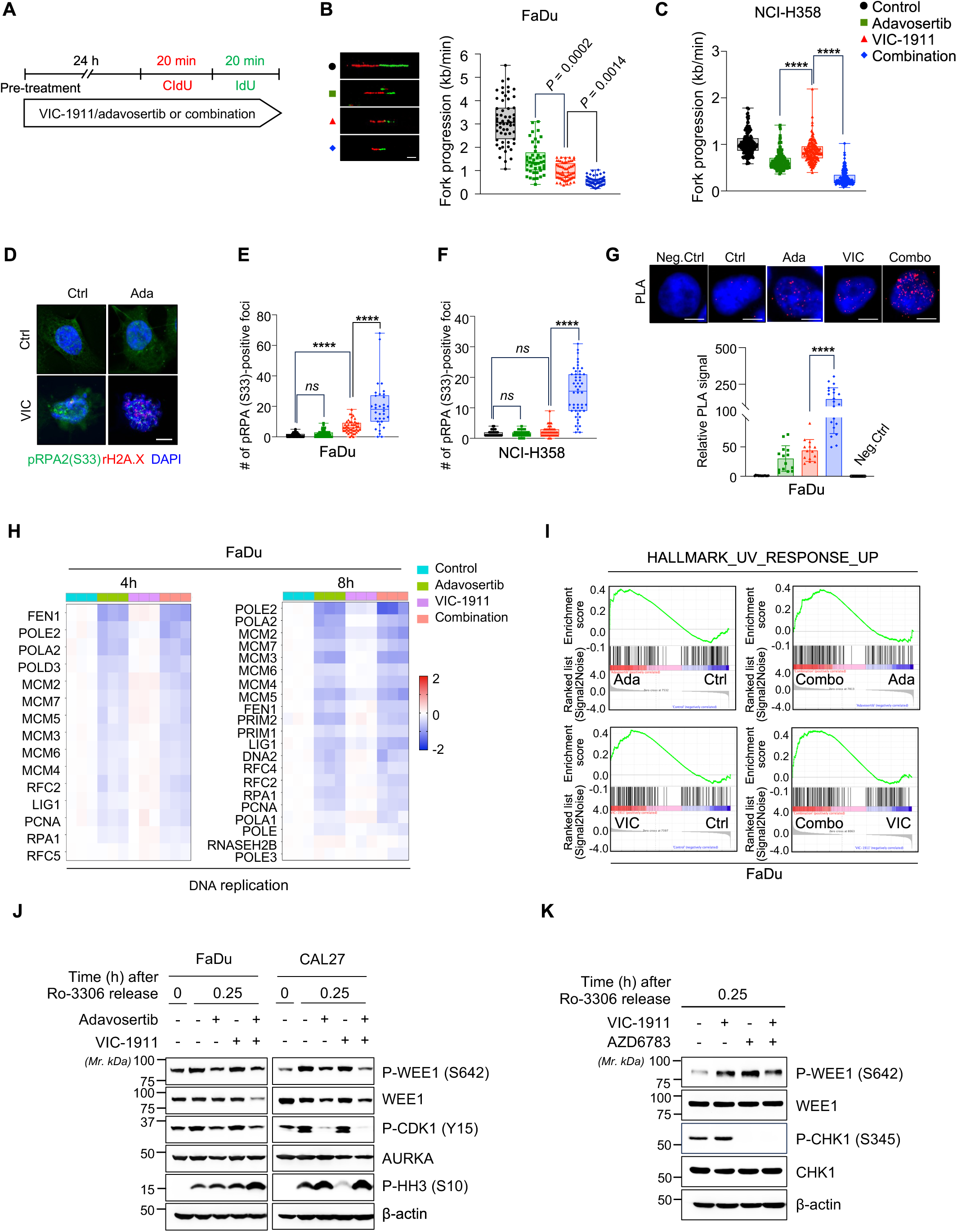
DNA replication stress induced by AURKA inhibition fosters WEE1 dependency. **A-C**, Fork progression speed in FaDu and NCI-H358 cells treated with control (Ctrl), adavosertib (Ada; 250 nmol/L), VIC-1911 (VIC; 125 nmol/L) or combination. **A**, The experimental design is shown. Cells were pre-treated with the indicated drugs for 24 hours and followed by sequential incubation with CIdU and IdU for 20 minutes each. **B,** Representative images of fibers (left) and quantification of fork progression as measured by fiber assays of FaDu cells (right). **C,** Quantification of fork progression as measured by fiber assays of NCI-H358 cells. Fibers (n=45-62 for FaDu; n=161-190 for NCI-H358) were examined. The box plots (as for those in all other figures) show median values (central lines) and all shown points as Min to Max. kb, kilobases. Scale bar = 10 μm. **D-F**, Synergistic increases of replication stress marker pRPA2 (S33)-positive foci in FaDu or NCI-H358 cells treated with VIC-1911/adavosertib. Cells were treated with control, adavosertib, VIC-1911 or combination for 24 hours and followed by immunofluorescence staining with pRPA2 (S33) and γH2A.X (S139) antibodies. Representative images of FaDu cells (**D**) and quantification of pRPA2 (S33)-positive foci in a cell of FaDu (**E**) and NCI-H358 (**F**). n ≥ 35 cells were examined. **G**, Transcription-replication conflicts induced by VIC-1911/adavosertib combination in FaDu cells. Cells treated with the indicated drugs as single or combination for 24 were assessed PLA intensity derived from interaction between RNA polymerase II and PCNA replisome. The negative control (Neg.Ctrl) shows primary antibodies only in cells treated with the combination. Representative images of PLA under different conditions, Scale bar = 10 μm (Top). Quantification of the PLA signals (Bottom). Each dot representative the mean PLA signal of all cells in a field as determined by area of PLA signal in total number of DAPI-positive nuclei, compared with the solvent control (Ctrl). n = 14 fields for the Neg.Ctrl, Ctrl, Ada and VIC; n = 21 fields for the Combo, and the data as shown mean ± SEM are representative of three independent experiments with consistent results. Statistical significance was determined by unpaired two-tailed *t*-test comparing two conditions (VIC-1911 vs Combination; \*\*\*\**P* < 0.0001). **H**, Heatmaps showing relative expression changes of DNA replication regulators genes under treatment with control, adavosertib, VIC-1911 or combination at different time points following by thymidine synchronization in FaDu cells. Only significantly changed genes based on RNA-seq analysis are shown. Each column indicates a biological replicate (n=3/condition). **I**, Pathway enhichment was dertermined by GSEA. **J,** Immunoblot analysis of the indicated proteins in FaDu and CAL27 cells upon the treatments at 15 minutes after CDK1 inhibitor-mediated synchronization. Cells were synchronized with Ro-3306 (9 μmol/L for 20 hours) and followed by release along with treatment of adavosertib and/or VIC-1911 versus control. Cell lysates were subjected to SDS-PAGE and immunoblotting with indicated antibodies. **K**, Immunoblot analysis of the indicated proteins in Ro-3306-mediated synchronized FaDu cells upon the treatments with VIC-1911 (125 nmol/L) and/or AZD6783 (1 μmol/L) for 15 minutes.

The DNA repair protein Replication Protein A2 (RPA2) is phosphorylated on serine 33 (S33) and binds to single-stranded DNA in the setting of replication stress and DNA damage and upregulates WEE1 to restrain cell cycle progression during DNA repair (35). AURKA inhibition augmented the number of pRPA (S33)-positive foci, in contrast to WEE1 inhibition had only a minimal effect on pRPA2 induction (Fig 3D-F). Increases of pRPA (S33)-positive foci were greatest in combination-treated cells, and combination treatment compared to control or single agents also resulted in higher intensity staining for γH2AX as a DNA damage marker, indicating higher replication stress and DNA damage (Fig. 3D-3F, Supplementary Fig S6E). Consistent with the hypothesis that inhibition of AURKA induces and that the activity of WEE1 is critical for its resolution

Transcription-replication conflicts (TRCs) are observed at regions of DNA replication fork stalling through formation of same-directional and converged replication forks when replication stress fails to elicit an appropriate replication stress response with cell cycle arrest; such TRCs exacerbate replication stress and lead to chromosomal instability. As a result, TRCs increase the interaction between the RNA polymerase II-transcription complex and proliferating cell nuclear antigen (PCNA) at the replication fork (36). We assessed TRCs in FaDu cells by proximity ligation assay (PLA) using antibodies against RNA polymerase II and PCNA and followed by quantification of PLA signal derived from all the cells in a microscopic field relative to control. We found a significant increase in PLA signaling upon combination AURKA and WEE1 inhibition; this was 200-fold higher than control, and more than 4-fold higher compared to AURKA or WEE1 inhibition alone (Fig. 3G), demonstrating that combined AURKA/WEE1 inhibition synergistically causes TRCs.

TRCs are also associated with increased formation of DNA-RNA hybrid structures called R-loops (36,37). To determine the effect of AURKA and WEE1 inhibition on R-loop formation, we conducted immunofluorescence staining using the S9.6 antibody that recognizes R-loops. WEE1 and AURKA inhibition independently resulted in a significant increase in R-loop content compared to control. Moreover, a larger increase in R-loop formation was observed upon AURKA/WEE1 co-inhibition relative to single agent treatment (Supplementary Fig. S6F). These data further substantiate the findings above indicating that AURKA and WEE1 inhibition in each case contributes to replication stress, providing the basis for synthetic lethality.

Stalled DNA replication should be reflected in the elevated expression of genes associated with late G1 and S phase. Using RNA sequencing, we identified elevated expression of mRNAs associated with DNA replication progression including *FEN1, POLE2, POLA1, MCM2, MCM7, MCM5*, *RPA1 and LIG1* either at 4h or 8 h after exposure to AURKA inhibitor compared to control (Fig. 3H). Consistent with stalled replication fork progression and elevated replication stress, in FaDu cells synchronized with double thymidine block and then released into adavosertib, VIC-1911, or the drug combination, multiple genes controlling DNA replication, including *POLE2, POLA2, MCM2, MCM7, MCM5, MCM3, MCM6* and *FEN1*, were downregulated at 4h and 8h after treatment with adavosertib or in combination with VIC-1911, suggesting stalled DNA replication progression (Fig. 3H). DNA damage response and DNA repair pathways are activated in response to DNA replication stress as recovery mechanisms (38). In this respect, our RNA-seq data showed that the UV-reponse gene set was enriched in combination VIC-1911/adavosertib-treated FaDu cells compared to control or those treated with single agents (Fig. 3I).

Given that WEE1 is activated to support an intact replication stress response, we hypothesized that AURKA inhibition-mediated replication stress induces WEE1 activation. To confirm induction of WEE1 activation, we treated FaDu cells synchronized by treatment with the CDK1 inhibitor Ro-3306 and found that at G_2_, determined by detection of phosphohistone H3 (Ser10) 15 minutes after release from synchronization, VIC-1911 treatment resulted in augmented WEE1 activity as shown by increased WEE1 phosphorylation at Ser 642 Activity of phosphorylated WEE1 was further confirmed by induction of the inhibitory Tyr 15 phosphorylation of CDK1. In contrast, combination VIC-1911/adavosertib decreased phosphorylation of WEE1 and CDK1 in HNSCC FaDu and CAL27 cells (Fig. 3J).

Because AURKA inhibition has been shown to activate ATR (39) and ATR is known to activate WEE1 (40), we hypothesized that ATR is critical to AURKA inhibitor-induced WEE1 activation. Therefore, we tested the effects of AURKA inhibition on WEE1 phosphorylation in the presence of the ATR inhibitor AZD6783. We found that phosphorylation of WEE1 by AURKA inhibition was prevented in the presence of AZD6783 (Fig. 3K). These data indicate that AURKA inhibition interferes with DNA replication fork progression to amplify replication stress, that the cytotoxicity of these effects is dampened by a replication stress response to which G_2_/M checkpoint arrest via WEE1 activation is integral, and that combined AURKA and WEE1 inhibition cooperatively enhances DNA replication stress while preventing the G_2_/M arrest that would allow repair of replication stress, resulting in cell death as cells proceed into M phase with unresolved replication stress.

### Triggering of mitotic catastrophe and cell death by combined Aurora kinase and WEE1 targeting is specific to AURKA inhibition

Combination of the AURKA inhibitor alisertib with a WEE1 inhibitor adavosertib is known to increase mitotic catastrophe and apoptotic cell death. In light of our finding that AURKA inhibition fosters WEE1 dependency through induction of replication stress, and because Aurora kinase B (AURKB) is upregulated in replication stress (41). [We sought to confirm a distinct role for selective AURKA inhibition in inducing replication stress. Since alisertib has been shown to have some degree inhibitory activity against AURKB when used at high doses (42). We tested to see whether We confirmed that the effects on mitotic regulation we had previously described with alisertib also result when the highly selective AURKA inhibitor VIC-1911 is used, ie. that these effects are not AURKB-mediated (Supplementary Fig. S7A and 7B). Therefore, we next assessed mitotic morphology 24 h after drug addition in control, adavosertib, VIC-1911 and combination-treated HNSCC FaDu and NSCLC NCI-H358 cells. Control and WEE1 inhibitor-treated cells showed a normal pattern of mitoses, whereas cells treated with VIC-1911 showed multipolar spindle formation and enlarged nuclei in both cell lines. Combination treatment caused profoundly defective mitoses with aggregated multipolar spindles, dispersed chromosome condensation and fragmentation, and micronuclei, consistent with mitosis-mediated cell death, also known as mitotic catastrophe (Fig. 4A; ref. 43). Furthermore, combined inhibition of AURKA and WEE1 significantly augmented the frequency of phospho-Histone H3 (S10)-positive cells with abnormal mitoses (Fig. 4B and 4C) compared to VIC-1911 single agent treatment.

**Figure 4.**
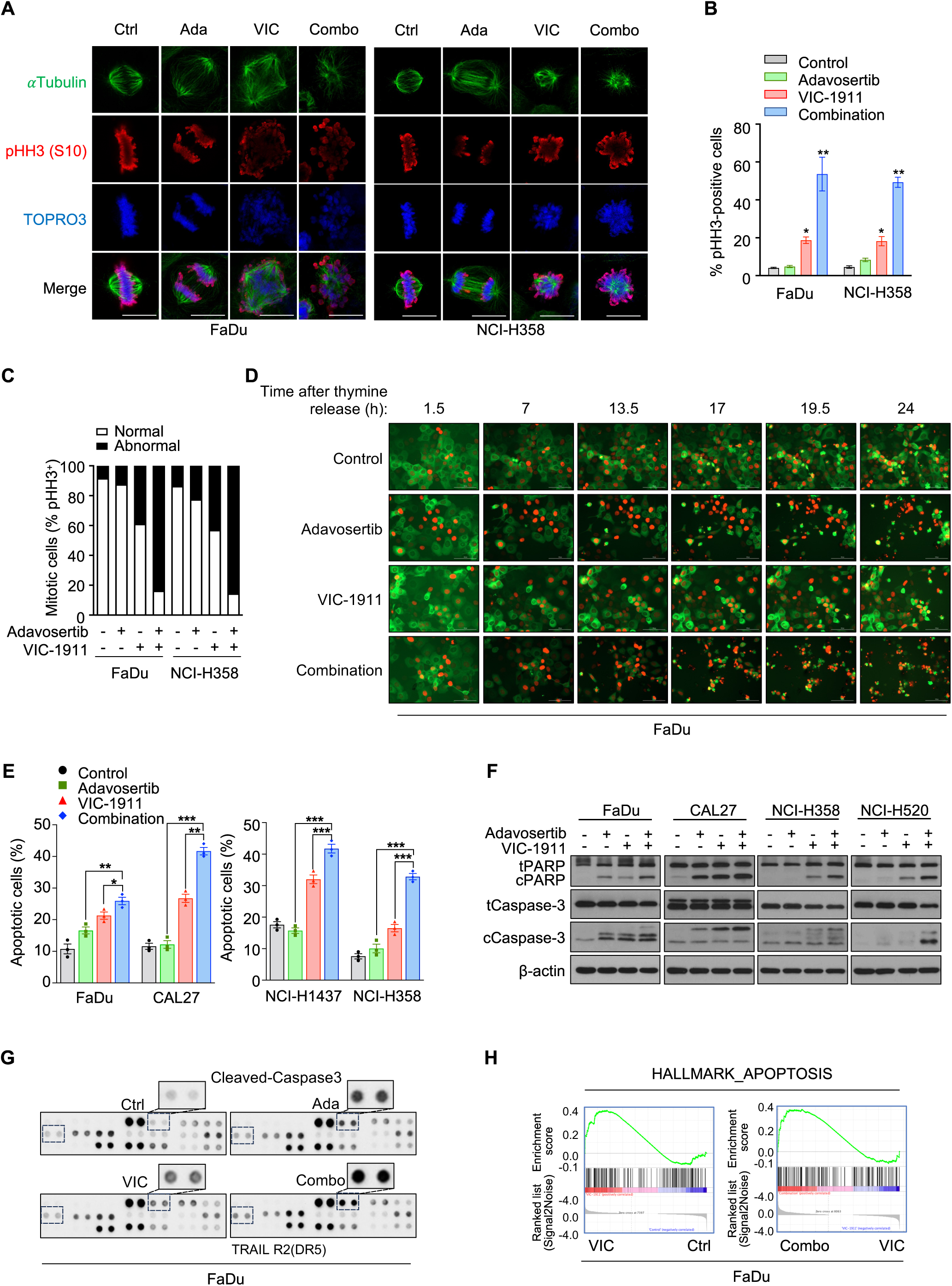
Concomitant treatment with VIC-1911 and adavosertib reveals synergistic inductions of defective mitoses and cell death in cancer cells. **A-C**, FaDu and NCI-H358 cells were treated with control, adavosertib (250 nmol/L), VIC-1911 (125 nmol/L) or combination for 24 hours, and followed by immunofluorescent staining with anti-tubulin (Green) and anti-phospho-Histone H3 (pHH3; S10; Red). Nucleus was stained with TOPRO3. **A**, Representative images of mitotic cells were captured by confocal microscopy. Scale bar = 10 μm. **B**, Percentage of dual pHH3 and TOPRO3-positive FaDu and NCI-H358 cells. Fields were randomly captured and counted more than 300 TOPRO3-positive cells. **C**, Percentage of normal or abnormal mitotic cells in pHH3-positive FaDu and NCI-H358 cells. **D**, Representative time-lapse live cell images of synchronized FaDu cells. EGFP-αTubulin and mCherry-Histone 2B stably expressing FaDu cells were synchronized by double thymidine block and followed by treatment with control, VIC-1911, adaovosertib or combination over 24 hours. Images were captured every 10 minutes interval and time indicates hours after release from double thymidine blocking with drug treatment. **E**, HNSCC (FaDu and CAL27) and lung cancer (NCI-H1437 and NCI-H358) cells were treated with the indicated single drug or combination for 48 hours and followed by staining with Annexin V and PI for apoptosis induction by flow cytometry analyses. Percentage of apoptotic cells in either Annexin V-positive or Annexin V/PI-positive cells. Each dot shows replicate from three independent experiments with similar results and data are shown as mean ± SEM. Statistical significance was determined by Student *t* test. \**P* < 0.05; \*\**P* <0.01; \*\*\**P* < 0.001. **F**, The indicated cell lines were exposed to the drugs singly or in combination for 48 hours. Cell lysates were subjected to SDS-PAGE and immuno-blotting with antibodies against cleaved PARP and cleaved Caspase-3 for apoptosis induction. **G**, Representative image of cleaved Caspase-3 induction in protein array for apoptosis panel of FaDu cells at 48 hours after exposure to the indicated drugs as single or combination. Enlarged signal are inserted. **H**, Pathway enhichment was dertermined by GSEA using RNA-Seq profile derived from FaDu cells treated with control, adavosertib, VIC-1911 or combination for 24 hours.

Although cells treated with VIC-1911 alone or in combination with adavosertib showed similar distributions in G_2_/M at 8h after release from double-thymidine synchronization, greater accumulation of cells in mitosis was observed following combination relative to VIC-1911 treatment, consistent with mitotic arrest (Supplementary Fig. S7A and S7B). Using time-lapse microscopy, we observed induction of mitotic catastrophe in FaDu cells stably expressing mCherry-H2B and EGFP-tubulin after treatment with control, adavosertib, VIC-1911 or combination. Mitotic cells were identified at an earlier time point - 7h after release from synchronization - under adavosertib treatment whereas VIC-1911 delayed mitotic entry and progress compared to control cells, with mitosis onset at approximately 17h after release; furthermore, abnormal mitoses in both adavosertib and VIC-1911 were observed at 19.5 h (Fig. 4D). Intriguingly, the proportion of severely defective mitoses harboring a mitotic catastrophe pattern of condensed/fragmented chromosomes and micronuclei drastically increased in FaDu cells treated with combination VIC-1911 and adavosertib, emerging at 17h (Fig. 4D).

Furthermore, the combination-treated cells exhibited a significant increase in apoptotic cell death, ranging from 25% to 42% relative to control or single drug treatment in both HNSCC and NSCLC cells, as determined by flow cytometry with Annexin V/PI staining (Fig. 4E and Supplementary Fig. S7C) or indicated by western blotting and protein apoptosis array for cleaved PARP and caspase-3 (Fig. 4F and 4G). We also observed enrichment of the hallmark_Apoptosis gene set in combination versus VIC-1911-treated FaDu cells (Fig. 4H), as well as upregulation of proapoptotic regulators including *DDIT3, BBC3, JUN, GADD45A, ATF4, CASP9, BCL2L11/BIM* and *PMAIP1/NOXA*, and downregulation of antiapoptotic regulators *BCL2, PIK3R2* and *PIK3CD* in both adavosertib and combination conditions versus control or VIC-1911 alone at 8h after release from thymidine-mediated synchronization (Supplementary Fig. S7D). Together, these data confirm that combined inhibition of AURKA and WEE1 cooperatively induces mitotic catastrophe and cell death in *TP53*-mutated cancer cells, across tumor types and in an AURKB-independent manner.

### The synergistic effect of AURKA/WEE1 co-inhibition is dependent on *TP53* status

Cyclin E (CCNE1) is expressed in replication stress and induces WEE1 expression; this marker of endogenous replication stress in cancer has been proposed as a molecular biomarker to select patients for WEE1 inhibitor therapy (30). We employed western blotting to see whether expression of CCNE1 is affected by AURKA or WEE1 inhibition in HNSCC cells. In FaDu cells synchronized by double thymidine block, VIC-1911 alone interestingly increased CCNE1 expression in particular, at 6 and 8h, and this CCNE1 level was further abolished by the addition of adavosertib (Fig. 5A). These data suggest that CCNE1 expression may not be necessary as a biomarker for clinical use of a WEE1 inhibition in combination with an AURKA inhibitor, as the AURKA inhibitor will induce the high replication stress state indicated by CCNE1 expression.

**Figure 5.**
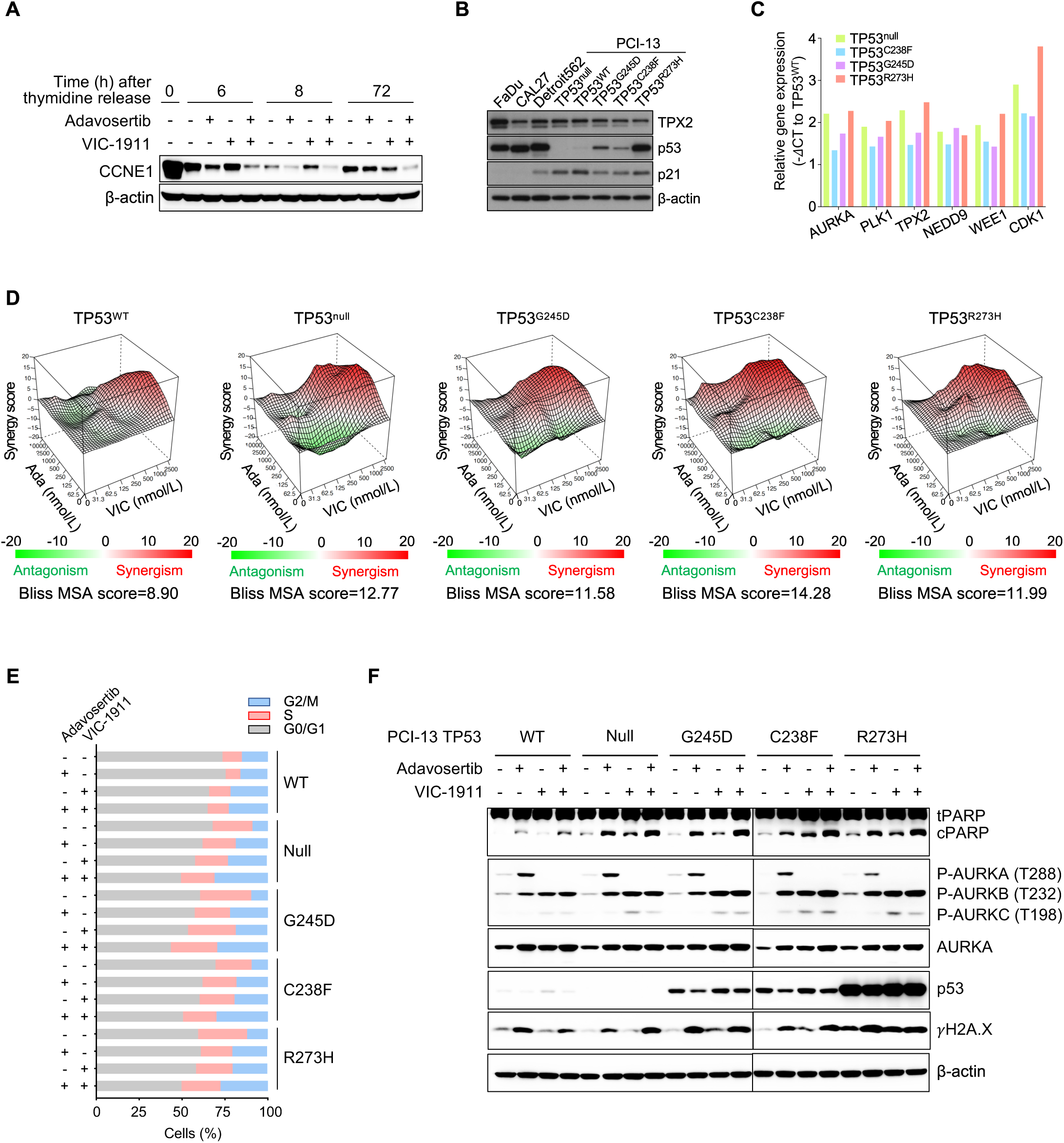
Combined AURKA/WEE1 inhibition attenuates CCNE1 expression and acts in TP53-dependent manner. **A**, FaDu and CAL27 cells were synchronized with Ro-3306 for 20 hours and released with complete media (supplement with 10% FBS) including the indicated drugs for 15 minutes. Cell lysates were subjected to SDS-PAGE and immuno-blotting. **B**, FaDu cells were synchronized with double thymidine block and followed by release with 10% FBS-containing media including control, VIC-1911 (125 nmol/L), adavosertib (250 nmol/L) or combination for the indicated time points. Cell lysates were subjected to SDS-PAGE and immuno-blotting. **C**, The indicated cell lines were cultured for 24 hours and cell lysates were subjected to SDS-PAGE and immuno-blotting for the indicated proteins. **D**, TP53- isogenic PCI-13 cells stably expressing control, *TP53 WT*, *G245D*, *C238F*, or *R273H* mutations were cultured in the complete media for 24 hours and RNA expression of the indicated genes were determined by qRT-PCR. mRNA expression of the genes was normalized with its expression in TP53 WT PCI-13 cells. **E**, Bliss synergy plotting for evaluating synergy score of combinational VIC-1911 and adavosertib treatment for 4 days. TP53-isogenic PCI-13 cells were treated with the indicated doses of adavosertib or VIC-1911 as single or combination as well as control for 4 days and cell viability was assessed by CellTiter-Glo assay. Loewe plotting were generated by conversion to bliss plots using cooperative correlation. Red area and Bliss MSA score indicate synergy; antagonism (< - 10), additive (-10 to 10) or synergism (> 10). **F**, Cell cycle distribution in synchronized PCI-13 TP53 isogeneic cells upon exposure to combinational treatment VIC-1911 (125 nmol/L)/adavosertib (250 nmol/L). Double-thymidine block-mediated synchronized PCI-13 cells were treated with control, adavosrtib, VIC-1911 or combination for 24 hours, and followed by flow cytometry of PI-stained cells G, PCI-13 cells were treated with control, VIC-1911 (125 nmol/L), adavosertib (250 nmol/L) or combination for 24 hours and cell lysates were subjected to SDS-PAGE and immuno-blotting.

Both AURKA and WEE1 were identified as potential targets in screens in the context of *TP53* mutation (2,4). To ascertain whether *TP53* mutation is necessary for the synergy observed here, we explored effects in cells isogenic for *TP53* mutations. We determined AURKA expression in PCI-13 isogenic cell lines of differing *TP53* genotypes including WT, null, *G245D*, *C238F* or *R273H*. We confirmed p53 expression in *TP53*-mutated HNSCC cells and PCI-13 *TP53* isogenic cell lines. *TP53* null PCI-13 cells showed completely deficiency of p53 expression and *TP53* WT cells modest expression of p53, consistent with the short half-life of WT p53, while cells bearing mutated *TP53* had higher p53 content. *TP53* WT PCI-13 cells expressed higher levels of the p53 effector p21, compared to other mutant cells (Fig. 5B). As expected, mRNA expression of *AURKA*, *PLK1*, *TPX2*, *NEDD9*, *WEE1* and *CDK1* genes was dramatically elevated in *TP53*-mutated PCI-13 isogenic cell lines relative to *TP53* WT cells (Fig. 5C).

We then assessed the impact of combination VIC-1911 and adavosertib treatment on cell viability of PCI-13 *TP53* isogenic cells. Synergy was observed in all cells harboring *TP53* mutations, whereas *TP53* WT cells showed only additive effect without synergy based on Bliss MSA scores (Fig. 5E). Intriguingly, synchronized *TP53* WT PCI-13 cells exhibited dramatically lower S (12.4%) and G_2_/M (22.8%) phase cell populations compared to *TP53* mutants including null (S, 19.6%; G_2_/M, 31.1%), G245D (S, 27.1%; G_2_/M, 29.9%), C238F (S, 19.5%; G_2_/M, 30.4%), R273H (S, 22.7%; G_2_/M, 27.6%)-harboring cells upon combination adavosertib and VIC-1911 treatment (Fig.5F and Supplementary Fig. S8). In agreement with our previous findings, we also confirmed that VIC-1911/adavosertib combination synergistically results in increases of cleaved PARP and γH2A.X in PCI-13 cells harboring *TP53* mutations and these increases were much greater in *TP53*-mutated compared with *TP53* WT PCI-13 cells (Fig. 5G). Thus, we conclude that *TP53*-mutated cells highly express AURKA and WEE1 and are more sensitive to combination treatment of VIC-1911 and adavosertib than are *TP53* WT HNSCC cells, suggesting the synergy may be dependent on *TP53* status.

### Combination VIC-1911 and adavosertib treatment leads to tumor regression and extended survival *in vivo*

We next determined the cooperative efficacy of VIC-1911 and adavosertib *in vivo* against various cancer cell-derived xenograft (CDX) and patient-derived xenograft (PDX) models. These included two HNSCC (FaDu and CAL27) and two NSCLC (NCI-H358 and A549) CDXs, the HNSCC PDXN04 (*MRE11*-mutated; *MRE11^E479G^*) and the LUAD PDXPRH (*TP53*-mutated; *TP53^R175H^*) derived from treatment-naïve patients. For CDXs, 4-5 mice were randomized to receive vehicle, adavosertib, VIC-1911 or combination treatment once tumors (n=6-8 per group) reached 150-350 mm^3^ prior to treatment initiation. Mice were administered vehicle, adavosertib (120 mg kg^-1^, *b.i.d*, daily), VIC-1911 (30 mg kg^-1^, 6-days-on/1-day-off) or combination by oral gavage for 3 weeks. Combination doses were initially determined across 3-point dose ranges in the FaDu xenograft model and the indicated doses were selected based on achieving synergistic tumor control with minimal toxicity. No significant differences in animal body weight change were detected between dose levels, but mild hepatotoxicity was noted at the highest adavosertib and VIC-1911 dose combination (Supplementary Fig. S8A-8C).

We observed that the humane endpoint was reached at 22 days after implantation in vehicle or VIC-1911-treated mice, consistent with our previous finding with alisertib in FaDu HNSCC xenograft models. Adavosertib treatment as a single agent exhibited modest effect in tumor growth, whereas combined VIC-1911/adavosertib treatment led to profound tumor control with stable disease, prolonged survival, smaller tumor size, and lower tumor weight compared to vehicle, VIC-1911 or adavosertib alone in FaDu CDXs (log-rank *P* = 0.0232; Fig. 6A, Supplementary Fig. S9A and S9B). No histopathological abnormalities of other organs including lung, intestine, kidney or spleen were observed (Supplementary Fig. S9C). We further confirmed that VIC-1911 plus adavosertib treatment enhanced tumor control in the HNSCC CAL27, NSCLC NCI-H358 and A549 CDXs models (Fig. 6B-6D, and Supplementary Fig. S10A-S10C). Combination-treated tumors were stable for a further 21 days after treatment was stopped in CAL27 CDXs, suggesting that the combination effect is durable (Fig. 6B).

**Figure 6.**
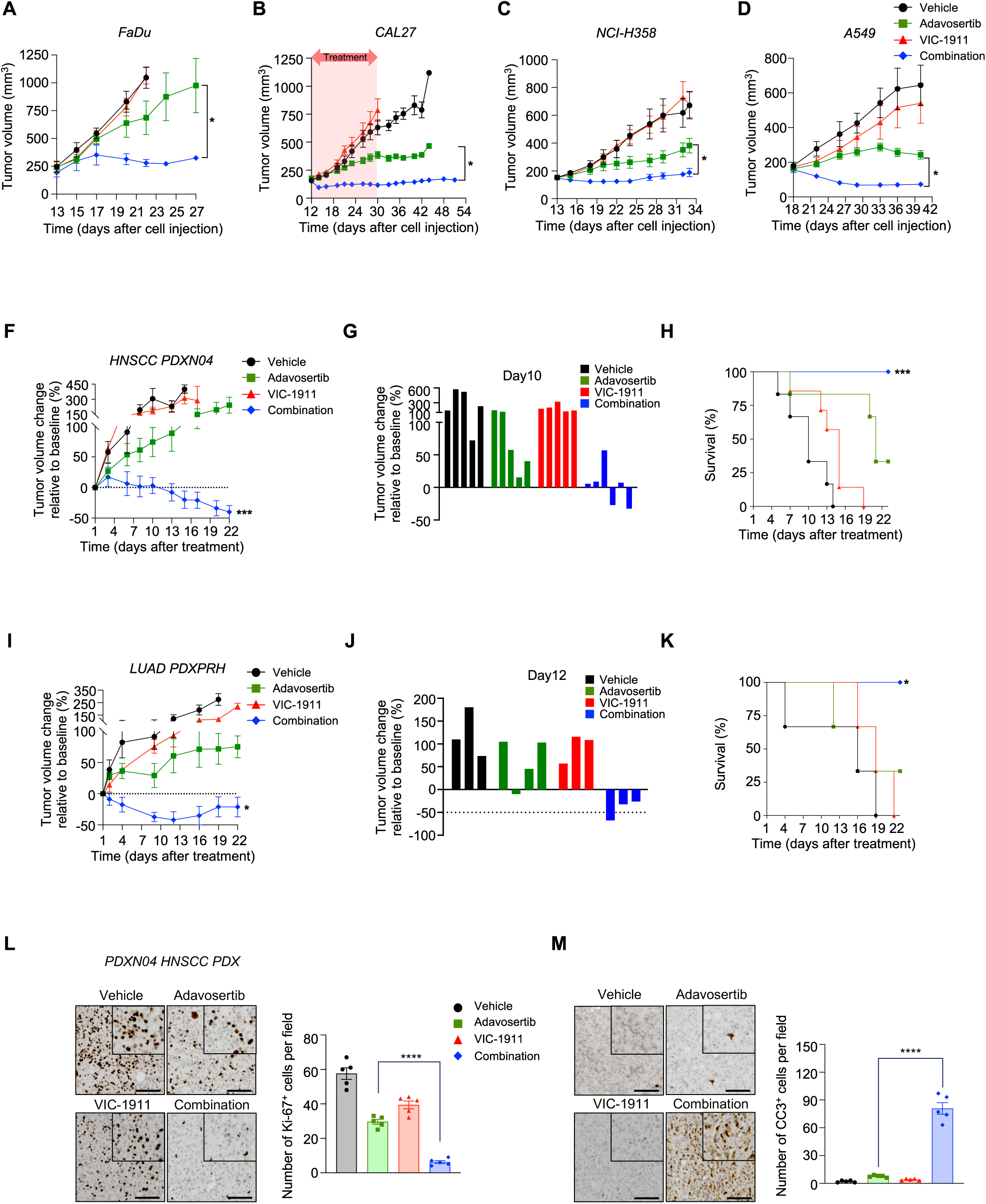
Therapeutic efficacy of combined AURKA/WEE1 inhibitor treatment in HNSCC and LUAD models *in vivo*. **A-D**, Mice bearing FaDu (**A**), CAL27 (**B**), NCI-H358 (**C**) or A549 (**D**) subcutaneous xenografts (FaDu n = 4; CAL27 n = 8; NCI-H358 n = 6; A549 n =5 per group) were treated daily (6 days on-1 day off) with vehicle, adavosertib 120 mg kg^-1^, VIC-1911 30 mg kg^-1^, or the combination for 3 weeks when tumor reached ∼200 mm^3^. Tumor growth curves were determined by measuring tumor volumes other day. **B**, CAL27 xenografts were treated as described above and tumor growth was further monitored after treatment duration was completed. Data are shown as mean ± SEM. *, *P* < 0.05. **F-H**. Mice bearing HNSCC PDX PDXN04 xenografts (n = 5-6) were treated as described above. **F**, Relative changes in tumor volume of HNSCC PDX model PDX04 during treatment with the indicated inhibitors. Relative tumor volume was calculated normalizing the measured tumor volume to the tumor volume on baseline (Day 1). **G**, Waterfall plot showing tumor volume changes after 10-day treatment. **H**, Kaplan-Meier survival curve of PDXN04 xenografted mice treated as indicated. Data are shown as mean ± SEM. ***, *P* < 0.001. **I-K**, Mice bearing LUAD PDX PDXPRH xenografts (n = 3-4) were treated as described above. **I**, Relative changes in tumor volume of HNSCC PDX model PDX04 during treatment with the indicated inhibitors calculated as described above. **J**, Waterfall plot showing tumor volume changes after 12-day treatment. **K**, Kaplan-Meier survival curve of PDXPRH xenografted mice treated as indicated. Data are shown as mean ± SEM. *, *P* < 0.05. **L** and **M**, Representative images of IHC staining for Ki-67 (L) and cleaved caspase-3 in tumor samples at the endpoint. Data are quantified in the graphs. Scale bars = 100 μm. Inset, high-power field. One-way ANOVA multiple comparison with Tukey’s correction were used to analyze the data. ****, *P* < 0.0001.

To enhance the relevance to human cancer therapy of the synergistic antitumor effects of combined AURKA and WEE1 inhibition we describe here, we next examined the *in vivo* efficacy of combination therapy in HNSCC (PDXN04; *MRE11*-mutated) or NSCLC (PDXPRH; *TP53*-mutated) PDXs. Tumor tissues reconstituted by passaging were implanted into dorsal flanks of NSG mice by surgical incision, and mice were randomly grouped as described above in CDX studies. Mice were treated with vehicle, adavosertib, VIC-1911 or combination by oral gavage for 6 days per week up to three weeks; tumor growth was assessed by change of tumor volume relative to baseline (due to greater baseline tumor size heterogeneity in PDX models). Whereas neither VIC-1911 nor adavosertib monotherapy prevented tumor progression of PDXN04 and PDXRPH PDXs, the combined VIC-1911/adavosertib treatment led to prominent tumor regression after 10 days of treatment and prolonged survival in the PDXN04 HNSCC (log-rank *P* < 0.0005; Fig 6F-6H) and PDXPRH NSCLC PDXs (log-rank *P* = 0.0172; Fig. 6I-6K), without weight loss (Supplementary Fig. S10D and S10E).

Correspondingly, there was marked induction of apoptotic cell death and elimination of proliferation as demonstrated by cleaved caspase-3 and Ki-67 staining, respectively, in combination-treated compared to either control- or single agent-treated tumors (Fig. 6L, 6M, Supplementary Fig. S11A and S11B). Tumoral micronuclei and multipolar spindles were detected in combination-treated HNSCC CAL27 CDX and PDXN04 models consistent with the previous observations in confocal and live cell imaging (Supplementary Fig. S11C and S11D). Immunohistochemistry confirmed that combined AURKA and WEE1 inhibition strongly led to increased R-loop formation and intensity of staining positive for phosphorylated γH2AX and 53BP1 in PDXN04 HNSCC tumors (Fig. 7A and 7B). Taken together, our data indicate that combined AURKA/WEE1 kinase inhibitor treatment results in notable antitumor effects with tumor regression in cell-line and patient-derived *in vivo* models of HNSCC and NSCLC, providing explicit preclinical confirmation of efficacy for this synthetic lethal therapeutic approach to eradicate tumors harboring *TP53* mutation.

**Figure 7.**
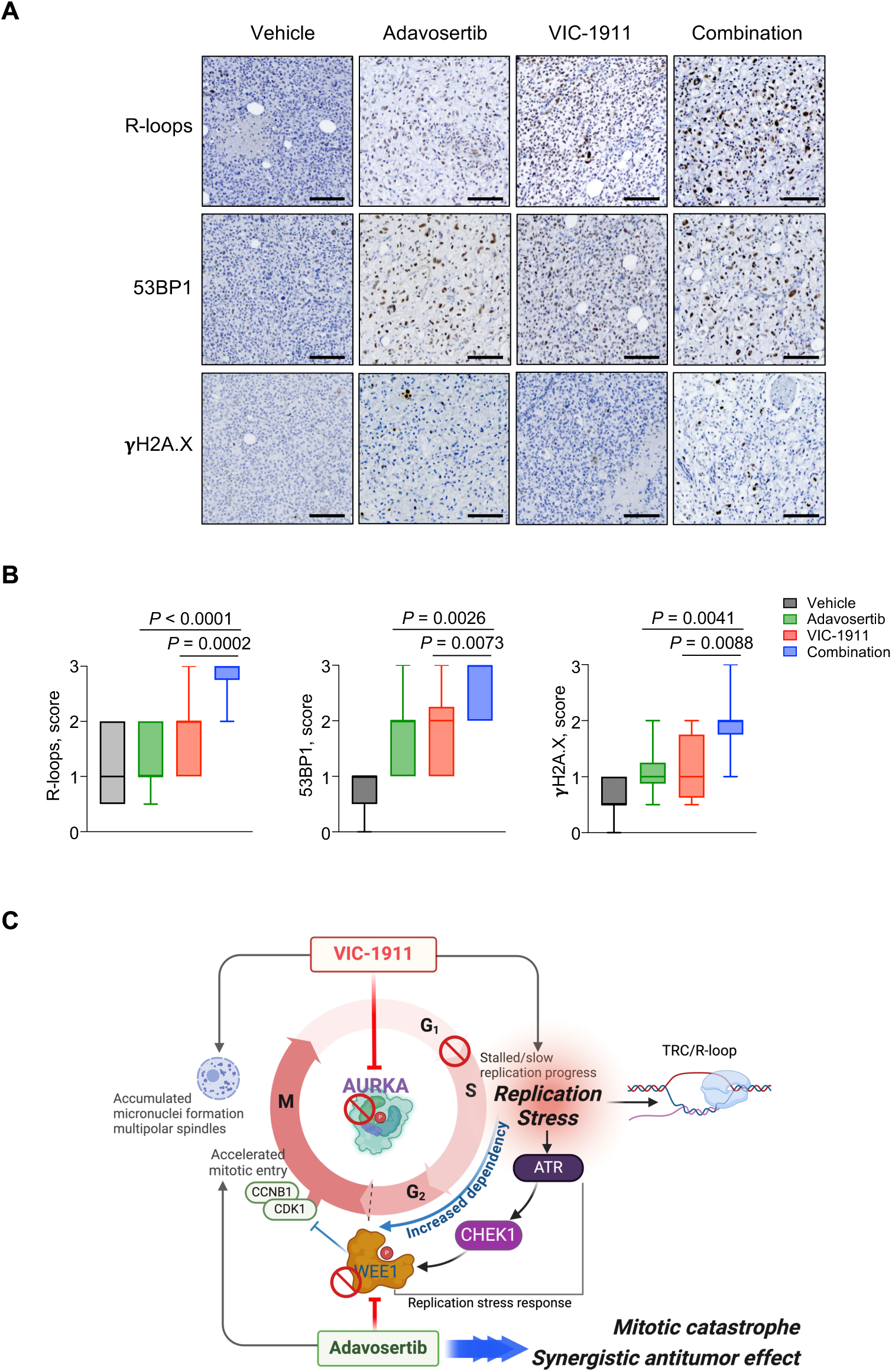
Concommitant AURKA/WEE1 inhibition increases R-loops and DNA damage in *in vivo*. **A,** Representative PDXN04 tumor sections showing R-loop and DNA damage markers 53BP1 and rH2A.X positivity. **B**, Box plots represents quantification of R-loop, 53BP1 and γH2A.X-positive intensity. Statistical significance was determined by two-way ANOVA using turkey comparison (n ≥10 sections from the animal described in A were evaluated). **C.** Schematic model summarizing the findings in this study.

## DISCUSSION

We here for the first time elucidate the mechanism of the striking synergy observed between AURKA and WEE1 inhibition in *TP53*-mutated carcinomas by interrogating the ways in which combination treatment with AURKA/WEE1 inhibitors impacts DNA replication stress and the normal replication stress response. Replication stress is broadly characterized by the slowing or stalling of DNA replication and is an important cause of the genome instability that is a hallmark of cancer (26). Faulty DNA replication progression is primarily repaired by a replication stress response dependent upon G_2_/M checkpoints; however, if repair fails, replication stress drives mitotic abnormalities including micronuclei formation, defective spindle assembly, and chromosome segregation errors which may be lethal to the cell (38,44,45). Such replication stress-driven aberrant mitoses result in mitotic catastrophe (43,46). It has previously been demonstrated that the AURKA/TPX2 complex contributes to replication fork stability and double-strand break repair by negatively regulating 53BP1, a process requiring AURKA catalytic activity (7). Here, one significant finding of our study is that AURKA inhibition by VIC-1911 dramatically impeded replication fork progression, increased localization of phosphorylated RPA2 to single-stranded DNA, and induced CCNE1 expression and the active, phosphorylated form of the mitotic checkpoint kinase WEE1. These effects foster a dependency on ATR/CHK1-mediated WEE1 activation for an intact replication stress response, a protective mechanism whose importance is particularly critical in cells with defective G_1_/S checkpoints, such as those with loss of p53 function (Fig. 7C). When WEE1 is inhibited in concert with AURKA, CDK1 and 2 are not phosphorylated and rather than the normal replication stress response with DNA repair, the cell loses restraint of mitotic entry. As AURKA inhibition has disrupted maturation of the spindle assembly complex, mitosis is defective, with high rates of mitotic catastrophe and apoptosis. We had previously observed that nuclear AURKA expression was particularly predictive of worse outcome in HNSCC (22), consistent with its importance in protecting DNA replication in *TP53*-mutated contexts. MRE11 plays roles in double-strand DNA damage repair and processing of replication stress as both an endonuclease/exonuclease and MRE11 has been proposed as a prognostic biomarker in HNSCC (47,48). Moreover, it has also demonstrated that a MRE11-mediated response to stalled replication forks can activate the cGAS-STING pathway, hence MRE11 is poised as a potential link between replication stress and inflammation (49). Of note, the HNSCC PDX model PDXN04 used in this study harbors a mutation (c.1435G>C; p.E479G) of the *MRE11* gene, thus further preclinical studies to evaluate *MRE11* mutation-mediated susceptibility to combination AURKA/WEE1 inhibitor treatment are warranted. Others have reported that AURKA-mediated regulation of DNA replication is a kinase-independent function (50,51); however, the catalytic domain-targeted AURKA inhibitor VIC-1911 clearly and dramatically impacts replication fork progression in our study. It would be worthwhile to determine whether VIC-1911 also affects noncatalytic functions of AURKA in this context.

We demonstrate the high efficacy of this approach with novel inhibitors of both AURKA and WEE1. Here, we adopted a newly synthetized highly selective AURKA inhibitor VIC-1911 (previously TAS-119) and, for the first time, confirmed the remarkable synergistic antitumor effect of combined VIC-1911 and adavosertib treatment across multiple solid tumors *in vivo* and in a set of patient-derived and cell line-derived models. Cumulatively, our studies suggest that WEE1 inhibition potentiates AURKA inhibitor therapy with promise to eradicate *TP53*-mutated cancers through aggravated replication stress and mitotic catastrophe.

We demonstrated that AURKA inhibition in combination with a WEE1 inhibitor synergistically reduced cell viability in cancer cells across 5 human solid tumors (Fig. 1 and supplementary Fig. S2-S5). Across a variety of different type of cancer cells, only the HPV-negative HNSCC cell line SCC61 failed to demonstrate synergy for combination adavosertib and VIC-1911 treatment (Fig. 1E). In this regard, we found that SCC61 cells harbor several gene mutations of interest that might be relevant to its lesser sensitivity to the combination, including the AURKA activator *AJUBA* (p.E310Q), and *CDC25C* (p.G224R; p.G254R; p.G263R; p.G297R).

It has been reported that AJUBA-mutant HNSCC cells express undetectable AJUBA expression and decreased PLK1 and Bora expression, resulting in resistance to inhibitors of PLK1, CHEK1/2 and WEE1 (52). In addition, CDC25C is phosphatase that dephosphorylates the CDK1/CCNB complex fostering accelated mitotic entry. Since these mutations are localized at the C-terminal of catalytic domain of CDC25C involved in phosphatase activity, loss of phosphatase function due to these mutations may be leave cells with poor mitotic entry, even in the presence of pharmacologic inhibition of the G_2_/M checkpoint kinases WEE1/PKMYT1 (53). This is consistent with the relative adavosertib resistance we observed, as well as with the possibility of accumulation in S phase with greater time for resolution of DNA damage. Future interrogation of these genes as determinants of AURKA-inhibitor induced replication stress would be of interest, and might yield clinically useful biomarkers for patient selection in trials of combined AURKA/WEE1 inhibition.

The synergistic antitumor effect of dual AURKA/WEE1 inhibition was further confirmed in cancer cells upon testing with the next generation WEE1 inhibitor azenosertib (Zn-c3) in combination with VIC-1911, indicating that this combination is unlikely due to off-target effects. *In vitro* experiments guided by confocal microscopy and live cell imaging, combination VIC-1911/adavosertib resulted in profoundly abnormal mitotic phenotypes including multipolar spindles, micronuclei formation and mitotic catastrophe, and induced marked apoptosis and an apoptotic gene signature in HNSCC and lung cancer cells. We further found remarkable *in vivo* pre-clinical evidence for this synergy in xenograft models, as shown by significant tumor regression and prolonged survival in PDXs of both HNSCC and lung cancer. These data suggest that combination AURKA/WEE1 inhibitor treatment could represent a highly active synthetic lethal strategy for treating *TP53*-mutated human cancers, historically a group of treatment resistant cancers for which targeted therapies have not been developed.

In this study, we explored potential biomarkers for future clinical trials. Although CCNE1 expression has been proposed as a patient selection marker for WEE1 inhibition (30), we observed that AURKA inhibition induces CCNE1 (consistent with our identification of its role in increasing replication stress), suggesting that baseline cyclin E expression and baseline high replication stress will not be required for clinical activity of this combination, since AURKA inhibition will induce this condition. Given the notion the negative regulation of AURKA by p53, we confirmed that AURKA/PLK1 expressions are highly elevated in most HNSCC cell lines harboring *TP53* mutations and PCI-13 isogeneic cells with different types of mutant TP53 compared to human normal cells and PCI-13 WT TP53 cells, respectively. In agreement with previous report demonstrating WEE1 as a critical therapeutic target in *TP53*-mutated cancers, we observed the increase of WEE1 expression in PCI-13 cells harboring *TP53* mutation. We show that synergy of combined AURKA/WEE1 inhibition was greatest when *TP53* was mutated, thus nominating *TP53* mutation as a suitable biomarker for this combination consistent with our pre-clinical evaluation. Taken together, our findings elucidate the mechanism underlying striking synergy for inhibition of the mitotic kinases AURKA and WEE1 through enhanced DNA replication stress and offer preclinical confirmation of efficacy for this synthetic lethal strategy for *TP53*-mutated carcinomas. Our data support the exploration of WEE1 inhibitors to enhance efficacy of AURKA inhibition in patients with *TP53*-mutated malignancies and warrant clinical investigation.

## METHODS

### Cell culture

Human HNSCC and lung cancer cell lines harboring *TP53* mutations were studied. HNSCC (FaDu, Detroit 562, CAL27, SCC25, and SCC-9), lung cancer (NCI-H520, NCI-H358, NCI-H1437, A549, NCI-H441, NCI-H23 and NCI-H1792), ovarian cancer (A2780 and SK-OV3), pancreatic cancer (MiaPaCa-2), hepatocellular carcinoma (HepG2), and normal human fibroblast BJ1 cell lines were purchased from the American Type Culture Collection (ATCC); HNSCC SCC61 cell line was purchased from Sigma-Aldrich; the UNC7 is a patient-derived cancer cell line (*kind gift of Wendell Yarbrough, University of North Carolina*) and PCI-13 isogenic cell lines of differing *TP53* including WT, null, G245D, C238F or R273H genotypes were kindly provided by Jeffrey Myers (MD Anderson Cancer Center). A normal human tracheobronchial epithelial cell line (NHTBE) was purchased from Lonza. FaDu, Detroit 562, CAL27, PCI-13 isogenic and BJ1 cells were maintained in EMEM media (ATCC), and SCC-9, SCC25, SCC9, SCC61 and UNC7 cells in DMEM/F12 media supplemented with 0.2 µg/mL hydrocortisone (Millipore-Sigma, H0135). All lung and ovarian cancer cell lines were maintained in RPMI-1640 media (Gibco) and all pancreatic cancer, and hepatocellular carcinoma cell lines in DMEM (Gibco). Media were supplemented with 10% fetal bovine serum and 1% Antibiotic-Antimycotic (Invitrogen). NHTBE cells were maintained in bronchial epithelial cell growth medium (BEGM) supplemented with BEGM bulletKit (Lonza; CC-3170). All cell lines were cultured under standard tissue culture conditions (5% CO_2_ at 37 °C) within less than 5 passages following resuscitation and routinely checked to be *mycoplasma* free using a MycoAlert mycoplasma detection kit (Lonza; LT07-418) and MycoStrip mycoplasma detection kit (Invivogen; rep-mys-10). All cell lines were authenticated using short tandem repeat DNA profiling (ATCC Human cell STR profiling service; 135-XV).

### Combination screening, clonogenic survival, soft agar and oncosphere formation assays

Cell viability, clonogenic survival, soft agar and oncosphere formation assays were performed as previously described (22). Cells were plated in 96-well plates and exposed to the indicated doses of VIC-1911 (dose range; 31.3-2,500 nmol/L), and adavosertib or azenosertib (dose range, 62.5– 10,000 nmol/L) in 8×8 combination format. Cells were incubated for four days in the presence of drugs to cover at least two doubling times. Cell viability was assessed with the CellTiter-Glo Luminescent Cell Viability Assay (Promega; G7572) to determine synergy by analysis of cooperativity screening, Loewe plotting and dose-response curves. For clonogenic survival assays, cells were plated in 12-well plates with low density and exposed to a single dose of VIC-1911 (125 or 250 nmol/L), adavosertib (250 or 500 nmol/L) or combination. Four days after treatment, cells were further incubated in the absence of drugs for an additional 10-14 days. Cells were stained with 0.1% crystal violet to evaluate clonogenic survival. For soft agar assays, cells were resuspended with 0.4% agarose-containing medium in 6-well plates contained bottom agarose and allowed to grow with exposure to the indicated drug(s) in three-dimensional conditions for four days. Colonies were further incubated in the absence of drugs, stained with p-iodonitrotetrazolium violet (Sigma-Aldrich; I10406) to select live colonies, and followed by analysis of the stained colonies using Image J software (NIH, USA). For oncosphere formation assays, cells were plated as single cell suspensions in serum-free media supplemented with 50 μg/ml Insulin (Millipore-Sigma; 11061-68-0), 20 μg/ml recombinant human EGF (PeproTech; AF-100-15), 10 μg/ml recombinant human basic FGF (PeproTech; 100-18B), 0.4% BSA (Millipore-Sigma; A1470), N-2 Plus Media Supplement (Thermo Fisher Scientific;17502048), B-27 Supplement (Thermo Fisher Scientific; 17504044) and 1% Antibiotic-Antimycotic in an ultra-low attachment plate (Fisher Scientific; 174927) for 7–9 days. The threshold for defining a colony or oncosphere was 10 cells, as indicated at >50 pixels in Image J or >100 μm on light microscopy.

The WEE1 inhibitors adavosertib (AZD1775; HY-10993) and azenosertib (Zn-c3; E1000) were purchased from MedChemExpress and Selleck Chemicals, respectively. The AURKA inhibitor VIC-1911 was obtained from Vitrac Therapeutics. All drugs were dissolved in Hybri-Max dimethyl sulfoxide (DMSO; Sigma-Aldrich; D2650) for *in vitro* experiments.

### Flow cytometry assays for cell cycle and apoptosis

Cell-cycle distribution and apoptotic cell death assays using flow cytometry were performed as previously described (22). Cells were treated with control, adavosertib (250 nmol/L), VIC-1911 (125 nmol/L) or in combination for 8 or 24 hours, following by double thymidine blocking. Cells were further stained with propidium iodide (PI)/RNase staining buffer (BD Pharmingen; 550825). For apoptosis assays, cells were exposed to control, adavosertib, VIC-1911 or in combination for 48 hours and followed by staining with Annexin V/PI staining buffer (BD Pharmingen; 556547). All stained cells were assessed on the BD LSRII Flow cytometer (BD Biosciences) and analyzed with FlowJo software (FlowJo LLC).

### Immunofluorescence staining and live cell imaging

Immunofluorescence staining was performed as previously described (22). For immunofluorescence staining, cells were seeded and treated with control, adavosertib, VIC-1911, or in combination for 24 hours. The treated cells were fixed with ice-cold methanol, permeabilized with ice-cold acetone, and blocked with 3% Bovine serum albumin (BSA) in PBS. Cells were further probed with primary antibodies overnight at 4°C, labeled with fluorescent-conjugated secondary antibodies (Alexa Fluor 488 and Alexa Fluor 647; Invitrogen; 1:1,000), and stained with TO-PRO-3 for nuclei and followed by mounting with ProLong Gold anti-fade reagent (Invitrogen; P36935). Stained cells were photographed with a Zeiss LSM710 confocal microscope (Carl Zeiss Microscopy, White Plains, NY, USA) at 100X magnification or an Echo Revolution microscope (Discover Echo, San Diego, CA, USA) at 10X-60X for representative images and quantitative analysis. The following primary antibodies were used: α-tubulin (Abcam; ab52866; 1:1,000), phospho-Histone H3 (S10; Cell Signaling Technology; #9701; 1,1,000), phospho-RPA32 (S33; Bethyl Laboratories; A300-246A; 1:400), phospho-Histone H2A.X (S139; Cell Signaling Technology; #9718; 1:500) and S9.6 (Sigma-Aldrich; MABE1095; 1:250). For live cell imaging, FaDu cells that stably expressed mCherry-H2B and EGFP-α-tubulin were synchronized with double thymidine block and released in the presence of control, adavosertib, VIC-1911 or combination treatment. Time-lapse microscopic images were captured every 5-8.5 minutes interval with 10X magnification (2x2 tiles) up to 24 hours using Cytation 5 cell imaging multimode reader (BioTek, Chalrotte, VT, USA). Representative images were deconvoluted using ImageJ and further processed in Adobe Photoshop 2024.

### Proximity ligation assay (PLA)

FaDu cells were plated in a glass bottom 96-well plate (Corning), treated for 24 hours with control, adavosertib (250 nmol/L), VIC-1911 (125 nmol/L), or combination, and fixed with methanol for 20 minutes. After permeabilization with ice-cold acetone for 1 minute, PLA were blocked with the Duo-link Blocking solution for 1 hour at 37 °C, followed by incubation with primary antibodies PCNA (Cell Signaling Technology; #13110; 1:1,000) and total RNAPolII (Santa Cruz Biotechnology; sc-55492; 1:1,000) overnight at 37 °C. Nuclei were counterstained using 4′,6- diamidino-2-phenylindole dihydrochloride (DAPI)-containing mounting medium. PLA were conducted using Duo-link PLA In Situ FarRed kit (Sigma-Aldrich; DUO92101) according to the manufacturer’s instructions. Images were captured with an Echo Revolution microscope at 20X for quantitative analysis and 60X for representative images; automatic position function was applied. Fourteen image fields per well were acquired for a minimum total of 800 cells per sample.

### DNA fiber assay

The DNA fiber assay was performed as described by Halliewell JA *et.al* (54). Briefly, cells were seeded in a 6-well plate and treated with control, adavosertib (250-500 nmol/L), VIC-1911 (125-250 nmol/L) or combination for 24 hours. Cells were treated with 5-chloro-2-deoxyuridine (CIdU; Sigma-Aldrich; C6891; 25 μmol/L) for 20 minutes and chased with 5-iodo-2-deoxyuridine (IdU; MedChemExpress; HY-B0307; 250 μmol/L) for an additional 20 minutes in the presence of the indicated drugs. Cells were lysed with spreading buffer (200 mmol/L Tris-HCl pH7.4, 50 mmol/L EDTA and 0.5% SDS) and DNA fibers spread on glass slides before fixation in a methanol:acetic acid (3:1) solution. After denaturation by 2.5 M hydrochloric acid, CIdU- and IdU-labeled fibers were stained with mouse anti-BrdU (BD Bioscience; B44; 1:500) and rat anti-BrdU (Abcam; ICR1; 1:500) antibodies, with Alexa Fluor 488-conjugated donkey anti-mouse IgG and Alexa Fluor 647-conjugated goat anti-rat IgG (Thermo Fisher Scientific; 1:1,000) as secondary antibodies. Stained DNA fibers were captured with an Echo Revolution microscope and analyzed with Image J to determine replication fork speed (kb/min). Statistical calculations were performed using GraphPad Prism 10 (GraphPad Software; Boston, MA, USA). An unpaired Student’s *t-*test was used to calculate statistical significance at *P* < 0.05.

### Immunoblotting and Protein array

Immunoblotting was performed as previously described (22). Whole cell lysates were prepared using RIPA lysis buffer (20 mmol/L Tris-HCl pH7.5, 150 mmol/L NaCl, 1 mmol/L EDTA, 1 mmol/L EGTA, 0.1% SDS, 1% sodium deoxycholate, 0.5% NP-40 and 0.5% Triton X-100) containing protease and phosphatase inhibitor cocktails (Roche). Protein concentrations were determined by Thermo Scientific™ Pierce™ BCA Protein Assay Kit (Thermo Fisher Scientific; PI23227) and clarified cell lysates were subjected to SDS-PAGE and immunoblotting using following the primary antibodies: cleaved PARP (D214; #5625; 1:1,000), PARP (#9542; 1:1,000), cleaved Caspase-3 (D175; #9664; 1:1,000), Caspase-3 (#9662; 1:1,000), phospho-AURKA/AURKB/AURKC (T288/T232/T198; #2914; 1:500), AURKA (#14475; 1:1,000), and WEE1 (#13084; 1:1,000) purchased from Cell Signaling Technology, Cycline E1 (Abcam; ab238081; 1:1000) and β-actin (Sigma-Aldrich; A2228; 1:10,000). Protein array was performed using the Proteome Profiler Human Apoptosis Array Kit (R&D systems; ARY009) according to the manufacturer’s instructions. Cell lysates were applied to antibody-conjugated array membranes, followed by incubation with HRP-conjugated streptavidin. Immunoblotting signals were visualized with a Chemiluminescent detection kit and captured using ChemiDoc MP (Bio-Rad; 12003154).

### Immunohistochemistry

Immunohistochemistry was performed as previously described (22). Collected tumor tissues were fixed in 10% formalin in PBS overnight and transferred to 70% ethanol. Tissues were further processed with paraffin-embedding and sectioning by Yale Pathology Tissue Service (YPTS). Sections were deparaffinized, re-hydrated and subjected to high-temperature antigen retrieval with 10 mM Sodium Citrate buffer (pH 6.0). Sections were blocked with BLOXALL endogenous blocking solution (Vector laboratories; SP-6000) and normal goat serum and incubated with primary antibodies against Ki-67 (MIB-1; Dako; M7240; 1:500) or cleaved capsase-3 (Cell signaling Technology; #9664; 1:500) overnight. Sections were washed and treated with SignalStain Boost IHC detection reagent (Cell signaling Technology; HRP; Rabbit; #8114 or Mouse; #8125). HRP-conjugated secondary antibodies were visualized by SignalStain DAB substrate kit solution (Cell signaling Technology; #8059) and followed by counterstaining with hematoxylin. The stained tissues were randomly captured using an Echo Revolution microscope (Echo a Bico company) with 10X automatic positioning and tiling.

### RNA sequencing and quantitative reverse transcription-PCR (qRT-PCR)

Total RNA was isolated from cells using miRNeasy mini kit (Qiagen; 217004) according to the manufacturer’s instruction, followed by determination of RNA quality by estimating the A260/A280 and A260/A230 ratios by Nanodrop. RNA samples were submitted to the Yale Center for Genome Analysis (YCGA) for RNA sequencing. RNA integrity was determined by running an Agilent Bioanalyzer gel, which measures the ratio of the ribosomal peaks. Samples with RNA Integrity Number (RIN) values of 7 or greater were advanced to library preparation. For library preparation, mRNA was purified from approximately 200ng of total RNA with oligo-dT beads and sheared by incubation at 94°C in the presence of Mg using a KAPA mRNA HyperPrep kit (Roche; KK8581). Following first-strand synthesis with random primers, second strand synthesis and A-tailing was performed with dUTP for generating strand-specific sequencing libraries. Adapter ligation with 3’ dTMP overhangs was ligated to library insert fragments. Library amplification was conducted to amplify fragments carrying the appropriate adapter sequences at both ends. Strands marked with dUTP were not amplified. Indexed libraries were quantified by qRT-PCR using a KAPA Library Quantification Kit (Roche; KK4854) and insert size distribution determined by the Agilent Bioanalyzer. Samples with a yield of ≥ 0.5 ng/μl and a size distribution of 150-300 bp were selected for further sequencing. For flow cell preparation and sequencing, sample concentrations were normalized and loaded onto an Illumina NovaSeq X Plus flow cell at a concentration that yielded 25 X 10^6^ passing filter clusters per sample. Samples were further sequenced using 101 bp paired-end sequencing on an Illumina NovaSeq X Plus according to the manufacturer’s instructions. The 10 bp unique dual index was acquired during additional sequencing reads automatically following completion of read 1. Data generated during sequencing runs were simultaneously transferred to the YCGA high-performance computing cluster. A positive control (prepared bacteriophage Phi X library) provided by Illumina was spiked into every lane at a concentration of 0.3% to monitor sequencing quality in real time.

For data analysis and storage, signal intensities were converted to individual base calls during a run using the system’s Real Time Analysis (RTA3) software. Base calls were transferred from the machine’s dedicated personal computer to the Yale High Performance Computing cluster for downstream analysis. Primary analysis - sample de-multiplexing and alignment to the human genome- was performed using Illumina’s CASAVA 1.8.2 software suite bcl2fastq2/2.20.0- GCCcore-10.2.0. Low quality reads were trimmed, and adaptor contamination removed using Trim Galore (v0.5.0). Trimmed reads were mapped to the human reference genome (hg38) using HISAT2 (v2.1.0) and gene expression levels were quantified using StringTie (v1.3.3b) with gene models (v27) from the GENCODE project. Differentially expressed genes were identified using DESeq2 (v 1.22.1; Ref. (55)). For qRT-PCR, RNA was extracted with miRNeasy mini kit (Qiagen) and followed by RT-PCR using iScript cDNA synthesis kit (Bio-rad) to synthesize cDNA for further qRT-PCR. Relative mRNA expression was measured by qRT-PCR using SsoAdvanced Universal SYBR Green Supermix (Bio-rad) and normalized to the expression of ACTB (β-actin).

### Animal studies

All mice were housed in a specific pathogen-free environment at the Yale University Animal Facility and treated in strict accordance with a protocol (#2022-11464) approved by the Yale Institutional Animal Care and Use Committee (IACUC). Six to eight-week-old female athymic nude (NU/J; 002019) and NSG (NOD.Cg-Prkdcscid Il2rgtm1Wjl/SzJ; 00557) mice purchased from The Jackson Laboratory were used in the cell-derived xenografts (CDX) and patient-derived xenografts (PDX) studies below, respectively. All mice were allowed access to sterile food and water *ad libitum*. For the CDX, one million cells (of the indicated HNSCC or lung cancer lines) were subcutaneously injected into the right flank of a nude mouse to establish a CDX model. Mice bearing CDX with a mean size of 150-300 mm^3^ received one of the following treatments: 1) vehicle; 2) adavosertib (120 mg kg^-1^, daily for 6 days, per os (p.o.)); 3) VIC-1911 (30 mg kg^-1^, daily for 6 days, p.o.); or 4) combination adavosertib (120 mg kg^-1^) + VIC-1911 (30 mg kg^-1^, daily for 6 days, p.o.) for three weeks. Agents were freshly prepared before dosing: adavosertib powder was dissolved with 5% DMSO, 40% Polyethylene glycol (PEG)-300 (Sigma-Aldrich; PHR3336), 5% Tween-80 (Sigma-Aldrich; 655207) and 50% PBS, and VIC-1911 powder was dissolved with filter-sterilized 0.5% (Hydroxypropyl) methyl cellulose (Sigma-Aldrich; 09963) in PBS. Briefly, for the PDX, tumor tissues were used within three passages of establishment from patients. Tissue fragments were rinsed in PBS and cut into 2-3 mm^3^ pieces with sterile razor blades, followed by implantation with Matrigel® Basement Membrane Matrix High Concentration (Corning; 47747-220). When tumor reached ∼300-550 mm^3^, mice were randomized and started on treatment with the above regime. Tumor volume was calculated as (*l* × *w*^2^) / 2, where *l* and *w* refer to the largest and smallest perpendicular dimensions at each measurement. Tumor-bearing mice were monitored daily for 6 days per week and mice were euthanized when tumor volume greater than 1,000 mm^3^ or tumor ulceration was observed, or 26 days after initial dosing, whichever came first. Tumor regression, median tumor volume, and treatment tolerability were also considered. Tumor weight was measured at the study endpoint. For toxicity profiling, whole blood and serum were collected from CDX animals at the endpoint for blood chemistry and diagnostic profiles with SuperChem/CBC (SA020; Antech Diagnostics).

### Calculation of drug parameters

For single-agent analysis, we estimated the IC_50_ using GraphPad Prism 10 with multiple dose-response models. For drug combinations, synergism was determined by the cooperative correlation, Loewe synergy and maximum synergistic area (MSA) scores and dose-response curve. we calculated both the Loewe synergy score and Chou-Talalay combination index. Loewe score was determined using the web-based SynergyFinder 3.0 (https://synergyfinder.fimm.fi) to define antagonistic (< -10), additive (-10 to 10) or synergistic (> 10) interactions (56). Chou-Talaly index was determined from the conjugated isobologram using CompuSyn software (https://www.combosyn.com) for additive effects (CI = 1), synergism (CI < 1), or antagonism (CI > 1; Ref. (57)).

### Statistical analysis

Plots and bar graphs depict the mean and standard error of the mean (SEM) as calculated by unpaired two-tailed *t*-test comparing two conditions or Student’s *t* test. Differences between treatment groups were determined by one- or two-way ANOVA followed by Bonferroni’s post-test. A Kaplan-Meier plot was generated to show survival with significance assessed by the log-rank (Mantel-Cox) test. Analyses were conducted using GraphPad Prism 10 and differences were considered to be significant at *P* < 0.05.

## Resource Availability

Further information and requests for resources and reagents should be directed to the lead contact, Barbara Burtness

## Authors’ Contributions

**J.W. Lee**: Conceptualization, data curation, resource, formal analysis, validation, investigation, visualization, methodology, writing-original draft, writing-review and editing. **J. Shi**: Formal analysis, validation, investigation, visualization. **J Barrantes**: Validation, investigation, visualization. **P. Chaurasia**: Formal analysis, investigation, visualization. writing-original draft. **S. Cruz-Gomez**: Investigation. **S. Kim**: Investigation, visualization. **C. Yang**: Formal analysis, Investigation, visualization. **J. Gu**: Formal analysis, methodology, data curation, visualization. **D. Zhao**: Formal analysis, methodology. **G. Kupfer**: Resource. **E. B. Perry**: Formal analysis. **J. P. Townsend**: Resource. **K. Politi**: Resource, **E. Golemis**: Resource, writing-review. **B. Burtness**: Conceptualization, resources, data curation, formal analysis, supervision, funding acquisition, writing-original draft, writing-review and editing.

## Acknowledgments

This work was supported by the Yale Head and Neck SPORE (NIH P50DE030707). We thank members of the Burtness lab for their valuable input. EG acknowledges support from NCI Core Grant P30 CA006927 (to Fox Chase Cancer Center).

