## Supplementary Figure S1 for "AURKA inhibition amplifies DNA replication stress to foster WEE1 kinase dependency and synergistic antitumor effects with WEE1 inhibition in cancers"

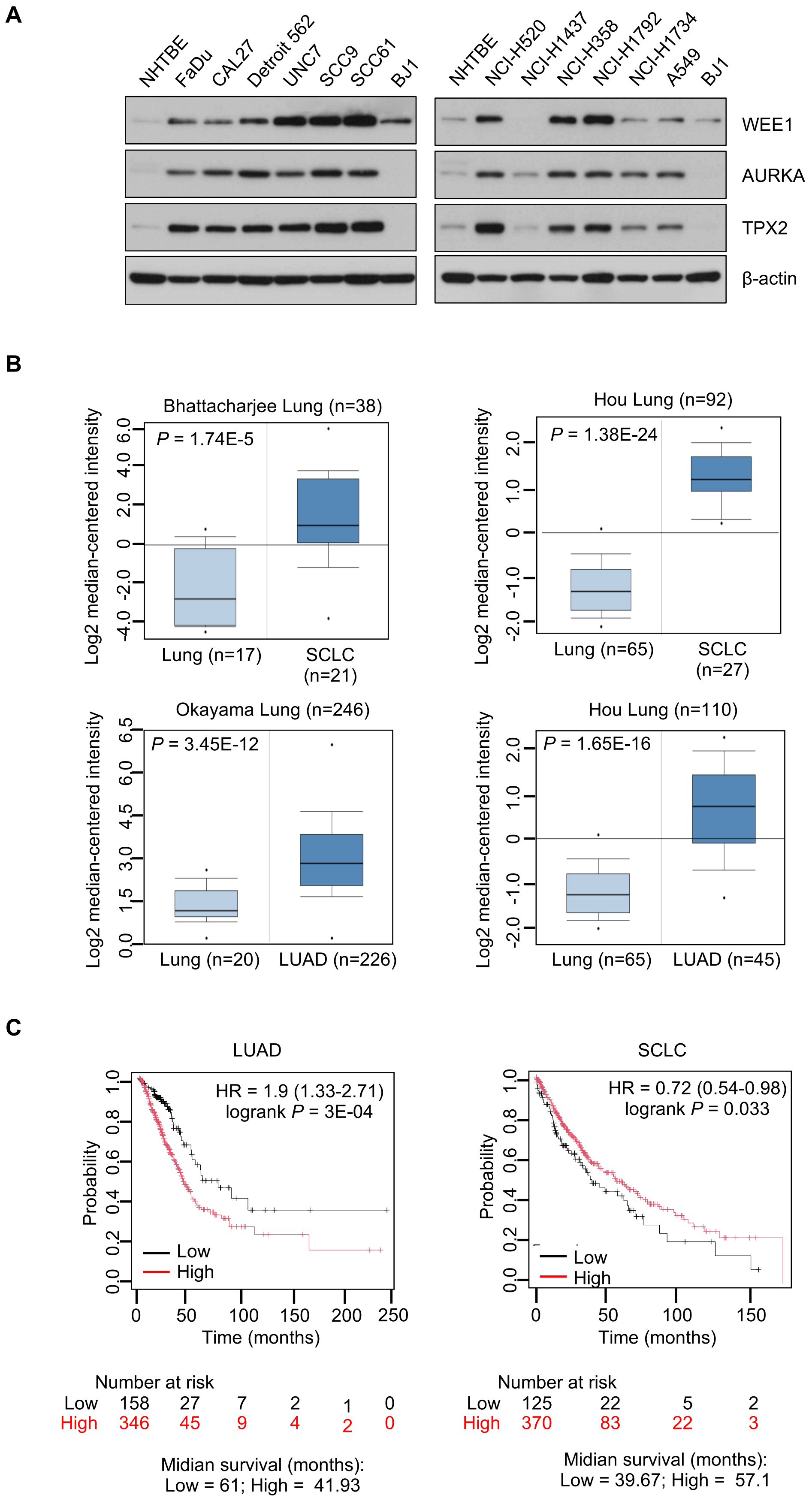


**Supplementary Figure S1**

**Supplementary Figure S1. AURKA is highly overexpressed in HNSCC and lung cancers and its expression is associated with poor outcome**. **A**, Cell lysates collected from the inidicated HNSCC/lung cancer cell lines were subjected to SDS-PAGE and immune blotting with antibody against WEE1, AURKA and TPX2. β-actin was served as loading control. **B**, AURKA expression was determined in NSCLC patients using oncomine database (Thermo Fisher Scientific) based on expression profiles (Bhattacharjee Lung n = 38; Hou Lung n = 202; Okayama Lung n = 246). AURKA expression in lung cancers was compared to normal counterparts and is showing statistically significant. **C**, Kaplan-Meier survival curve of lung cancer patients harboring AURKA overexpression in pan-cancer RNA-Seq database using km-plotter (http://www. kmplot.com).
