## Supplementary Figure S2 for "AURKA inhibition amplifies DNA replication stress to foster WEE1 kinase dependency and synergistic antitumor effects with WEE1 inhibition in cancers"

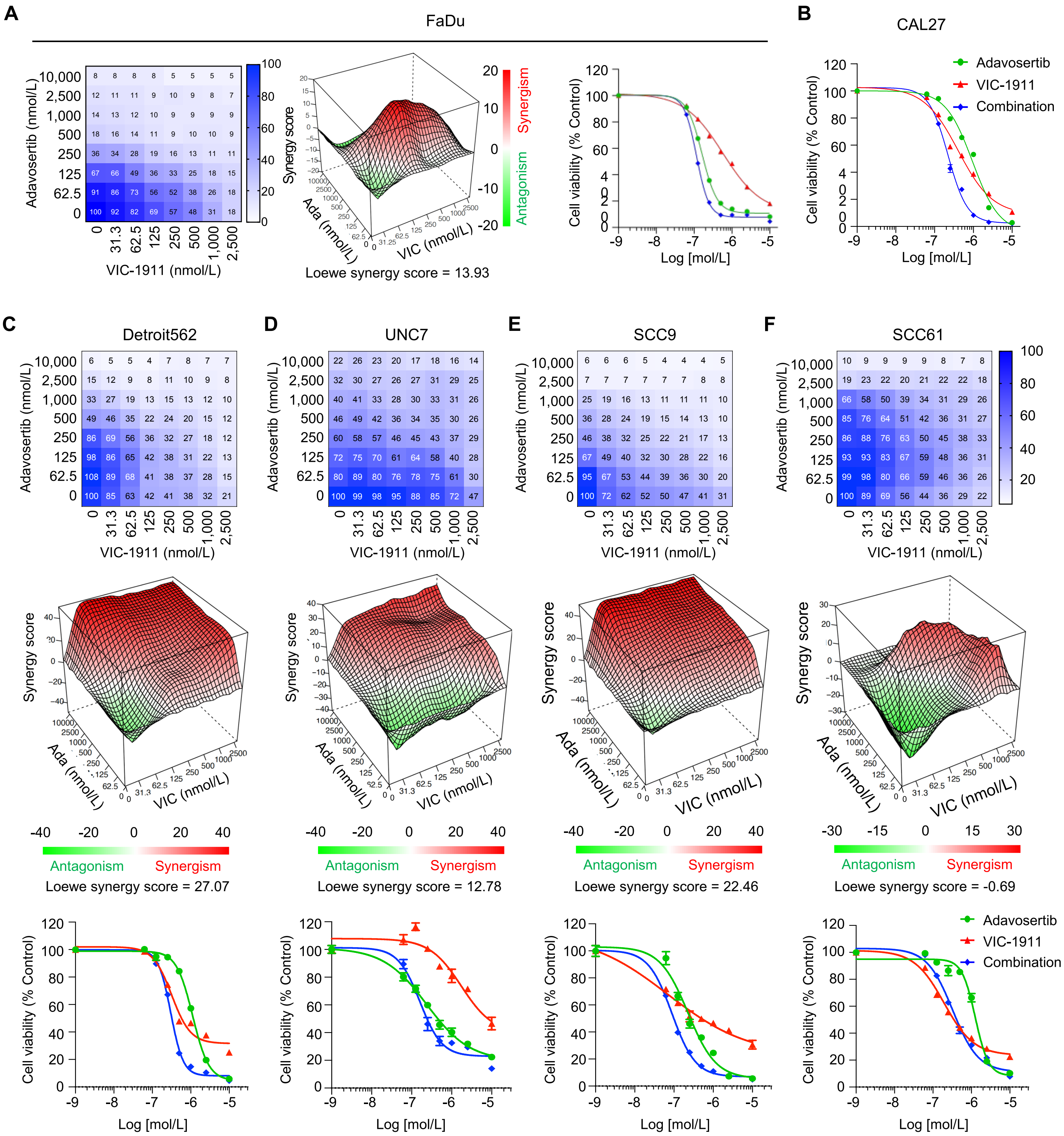


**Supplementary Figure S2**

**Supplementary Figure S2. Combination VIC-1911/adavosertib drug testing in extended HNSCC cell lines.** Human HPV-negative HNSCC cell lines including FaDu (**A**) , CAL27 (**B**; for Fig. 1C), Detroit 562 (**C**), UNC7 (**D**), SCC9 (**E**), and SCC61 (**F**) were treated with control, adavosertib (Ada; 0.0625-10 μmol/L), VIC-1911 (VIC; 0.0313-2.5 μmol/L), or 8 × 8 - combination for 96h. Cell viability was assessed by CellTiter-Glo assay and normalized to control treated cells for Fig. 1E. Number indicates % of cell viability related to control. Synergy of the combination was determined by cooperative correlation charts (A: Left; C-F: Upper), Loewe synergy score (A and C-F: Middle) and dose-response curves (A: Right; B for Fig. 1C; C-F: Bottom). Shown are the means of technical triplicates from one experiment and the data as shown mean ± SEM are representative of three independent experiments with consistent results.
