## Supplementary Figure S3 for "AURKA inhibition amplifies DNA replication stress to foster WEE1 kinase dependency and synergistic antitumor effects with WEE1 inhibition in cancers"

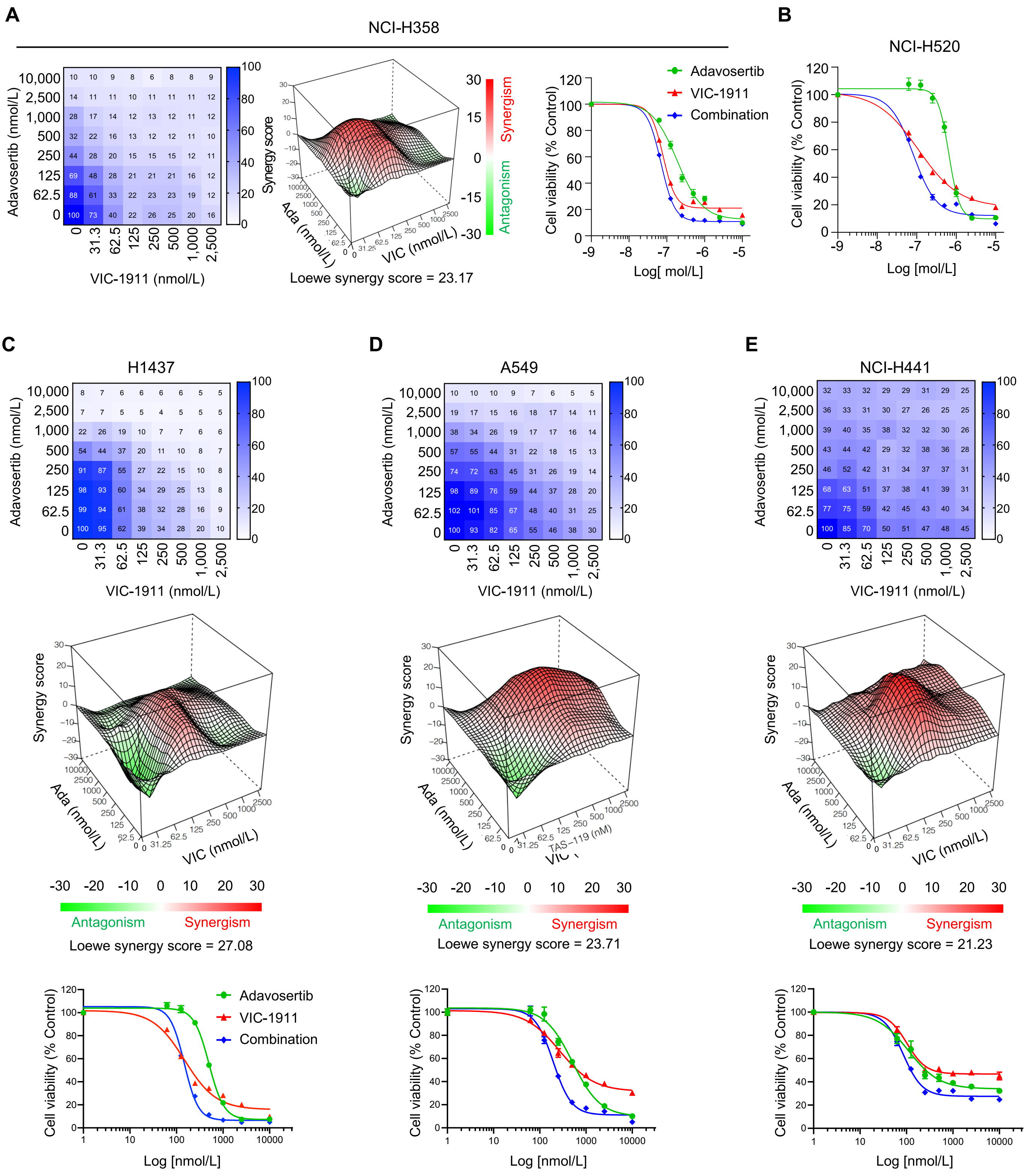


**Supplementary Figure S3**

**Supplementary Figure S3. Combination VIC-1911/adavosertib drug testing in extended NSCLC cell lines.** Human NSCLC cell lines including NCI-H358 (**A**) , NCI-H520 (**B**; for Fig. 1D), NCI-H1437 (**C**), A549 (**D**), and NCI-H441 (**E**) were treated with control, adavosertib (Ada; 0.0625-10 μmol/L), VIC-1911 (VIC; 0.0313-2.5 μmol/L), or 8 × 8 - combination for 96h. Cell viability was assessed by CellTiter-Glo assay and normalized to control treated cells for Fig. 1E. Number indicates % of cell viability related to control. Synergy of the combination was determined by cooperative correlation charts (A: Left; C-E: Upper), Loewe synergy score (A and C-E: Middle), and dose-response curves (A: Right; B for Fig. 1D; C-E: Bottom). Shown are the means of technical triplicates from one experiment and the data as shown mean ± SEM are representative of three independent experiments with consistent results.
