## Supplementary Figure S4 for "AURKA inhibition amplifies DNA replication stress to foster WEE1 kinase dependency and synergistic antitumor effects with WEE1 inhibition in cancers"

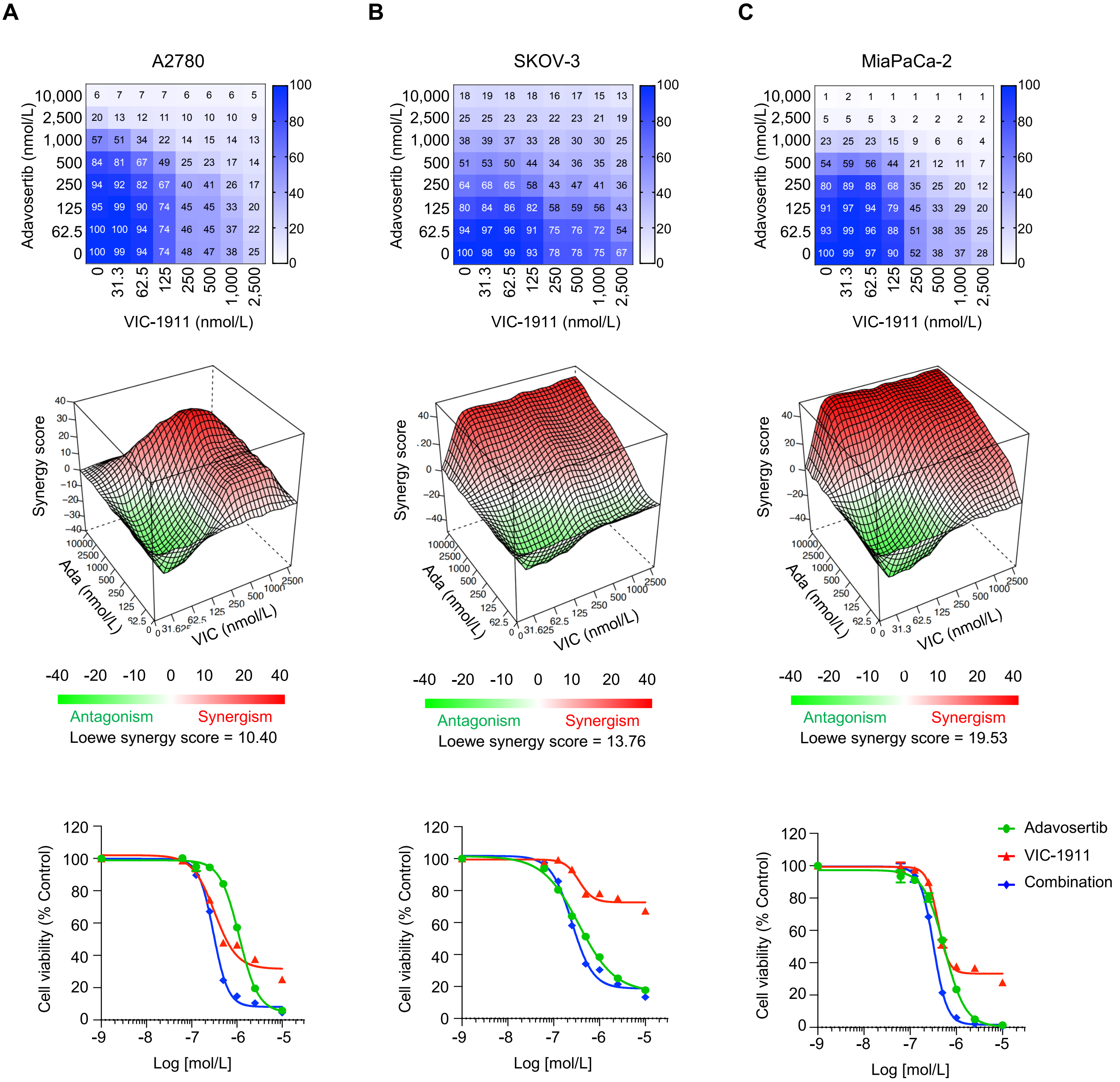


**Supplementary Figure S4**

**Supplementary Figure S4. Combination VIC-1911/adavosertib drug testing in various type of cancer cell lines.** Human cancer cell lines including ovarian cancer A2780 (**A**) and SKOV-3 (**B**), pancreatic cancer MiaPaCa-2 (**C**), and HCC HepG2 (**D**) were treated with control, adavosertib (Ada; 0.0625-10 μmol/L), VIC-1911 (VIC; 0.0313-2.5 μmol/L), or 8 × 8 - combination for 96h. Cell viability was assessed by CellTiter-Glo assay and normalized to control treated cells for Fig. 1E. Number indicates % of cell viability related to control. Synergy of the combination was determined by cooperative correlation charts (Upper), Loewe synergy score (Middle), and dose-response curves (Bottom). Shown are the means of technical triplicates from one experiment and the data as shown mean ± SEM are representative of three independent experiments with consistent results.
