## Supplementary Figure S5 for "AURKA inhibition amplifies DNA replication stress to foster WEE1 kinase dependency and synergistic antitumor effects with WEE1 inhibition in cancers"

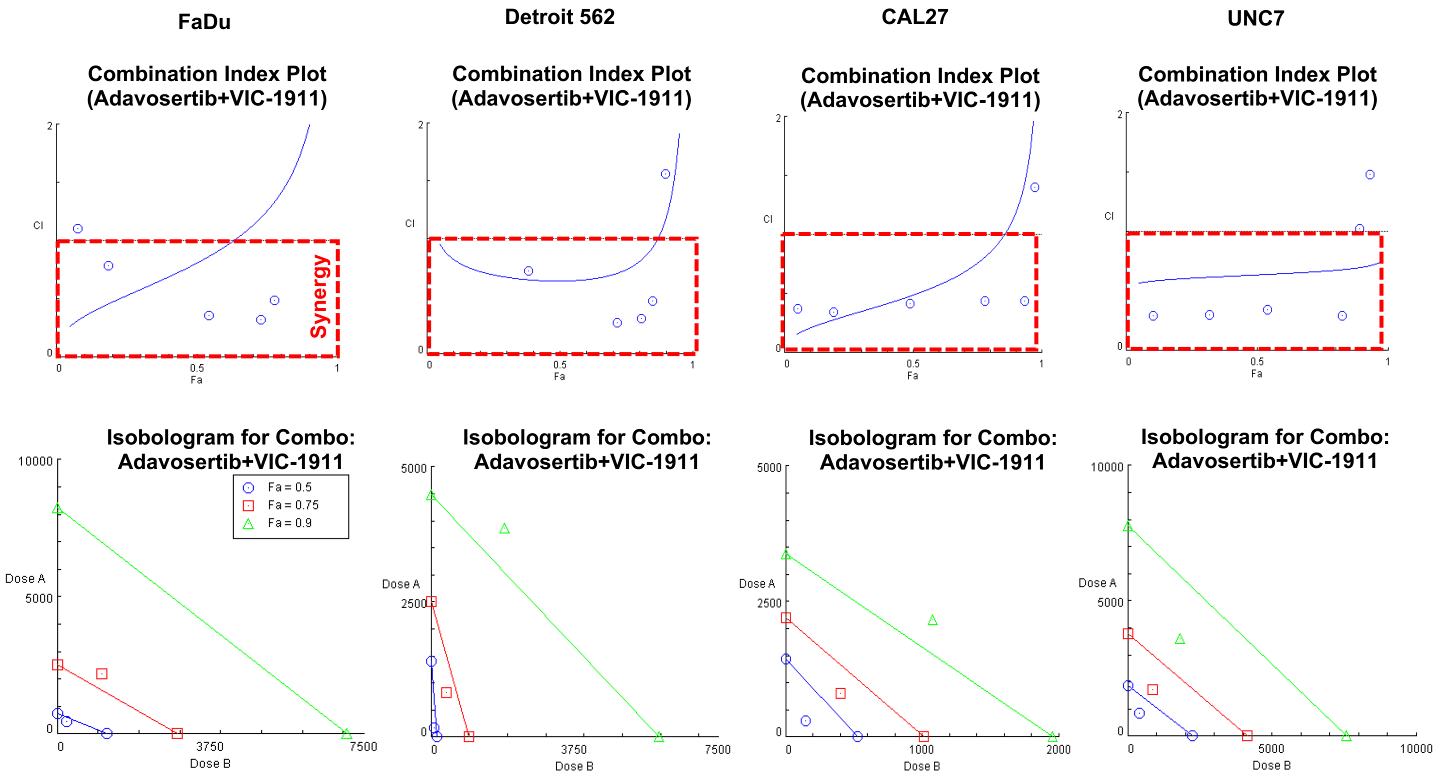


**Supplementary Figure S5. Synergy of combinational VIC-1911 and VIC-1911 treatment in HNSCC cells.** Cells were treated with control, adavosertib (50-10,000 nmol/L), VIC-1911 (25–5,000 nmol/L), or combination pairs and followed by cell viability assessed by CellTiter-Glo assay. Viable cells normalized by control treated cells were applied to the CompuSyn software (http://www.combosyn.com) to determine synergism based on Chou-Talalay combination index plot and isobologram. Red box in combination index plot and lower left of the hypotenuse indicate synergy (Fa: effective level; 0.5 (50%), 0.75 (75%), 0.9 (90%) synergy).
