## Supplementary Figure S6 for "AURKA inhibition amplifies DNA replication stress to foster WEE1 kinase dependency and synergistic antitumor effects with WEE1 inhibition in cancers"

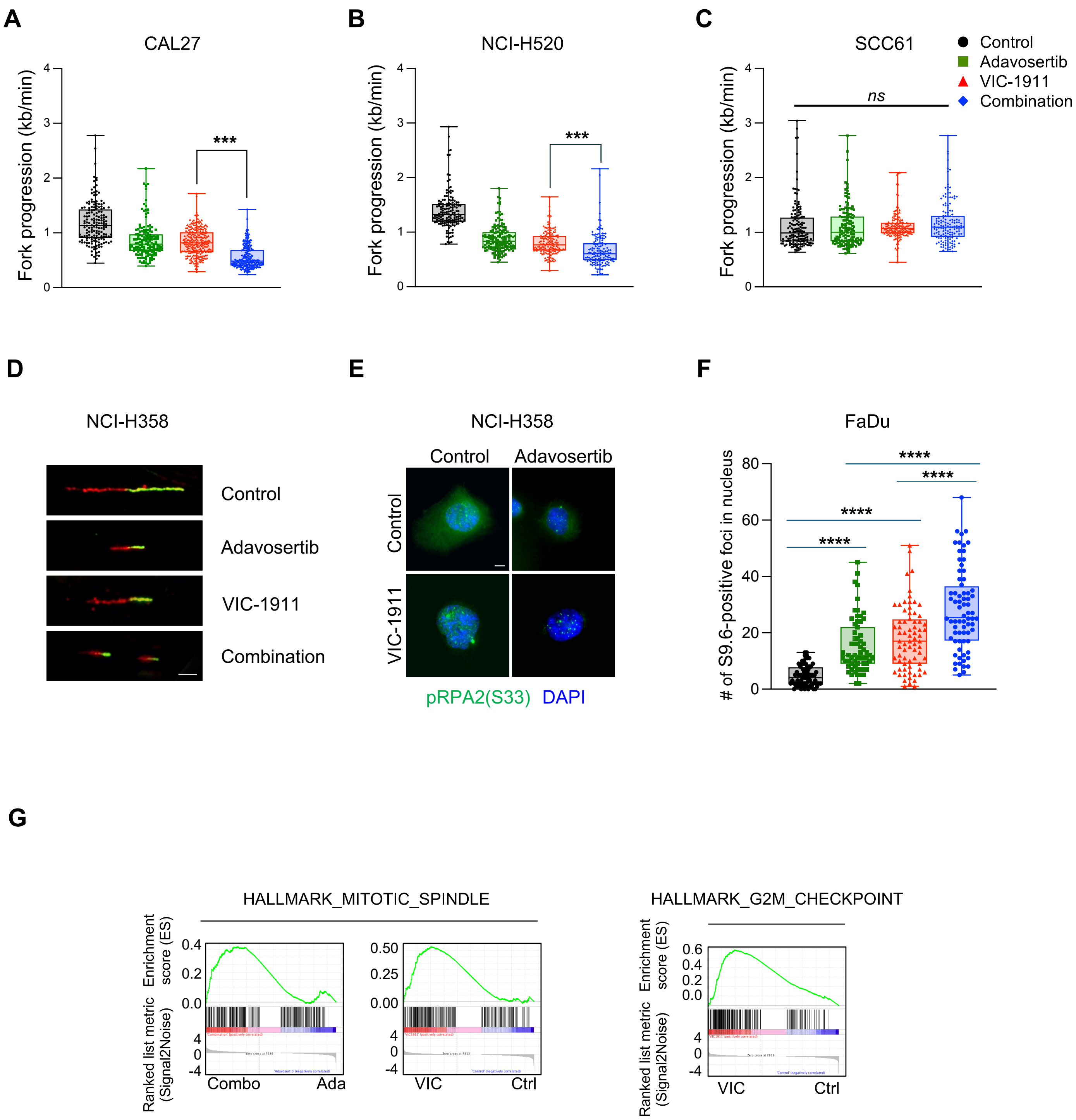


**Supplementary Figure S6**

**Supplementary Figure S6. DNA replication stress induced by VIC-1911/adavosertib combination in HNSCC and lung cancer *in vitro* models. A-C**, Fork progression speed in CAL27 (**A**), NCI-H520 (**B**) and SCC61 (**C**) cells treated with control, adavosertib (250 nmol/L), VIC-1911 (125 nmol/L) or combination for 24h and followed by fiber assay as described in the Methods. Quantification of fork progression as measured by fiber assays of the indicated cell lines. Fibers (n = 150-250 for CAL27; n = 150 for NCI-H520; n = 150 for SCC61) were examined. The box plots (as for those in all other figures) show median values (central lines) and all shown points as Min to Max. kb, kilobases. ***$P$ < 0.001. ns, non-significance. **D**, Representative images of NCI-H358 fibers in each treatment. Scale bar = 10 μm. **E**, Synergistic increases of replication stress marker pRPA2 (S33)-positive foci in NCI-H358 cells treated with VIC-1911/adavosertib. Representative images of NCI-H358 cells treated with control, adavosertib, VIC-1911 or combination for 24 hours and followed by immunofluorescence staining with pRPA2 (S33) antibodies and DAPI for nucleus staining. n ≥ 35 cells were examined. **G,** Gene set enrichment analysis (GSEA) of differential expression in FaDu cells upon the treatments. RNA-Seq data derived from FaDu cells treated with VIC-1911/adavosertib. FaDu cells were treated with control (Ctrl), adavosertib (Ada), VIC-1911 (VIC), combination (Combo) for 24h and followed by extraction of total RNA for further RNA sequencing. Pathway enrichment was determined by GSEA.
