## Supplementary Figure S7 for "AURKA inhibition amplifies DNA replication stress to foster WEE1 kinase dependency and synergistic antitumor effects with WEE1 inhibition in cancers"

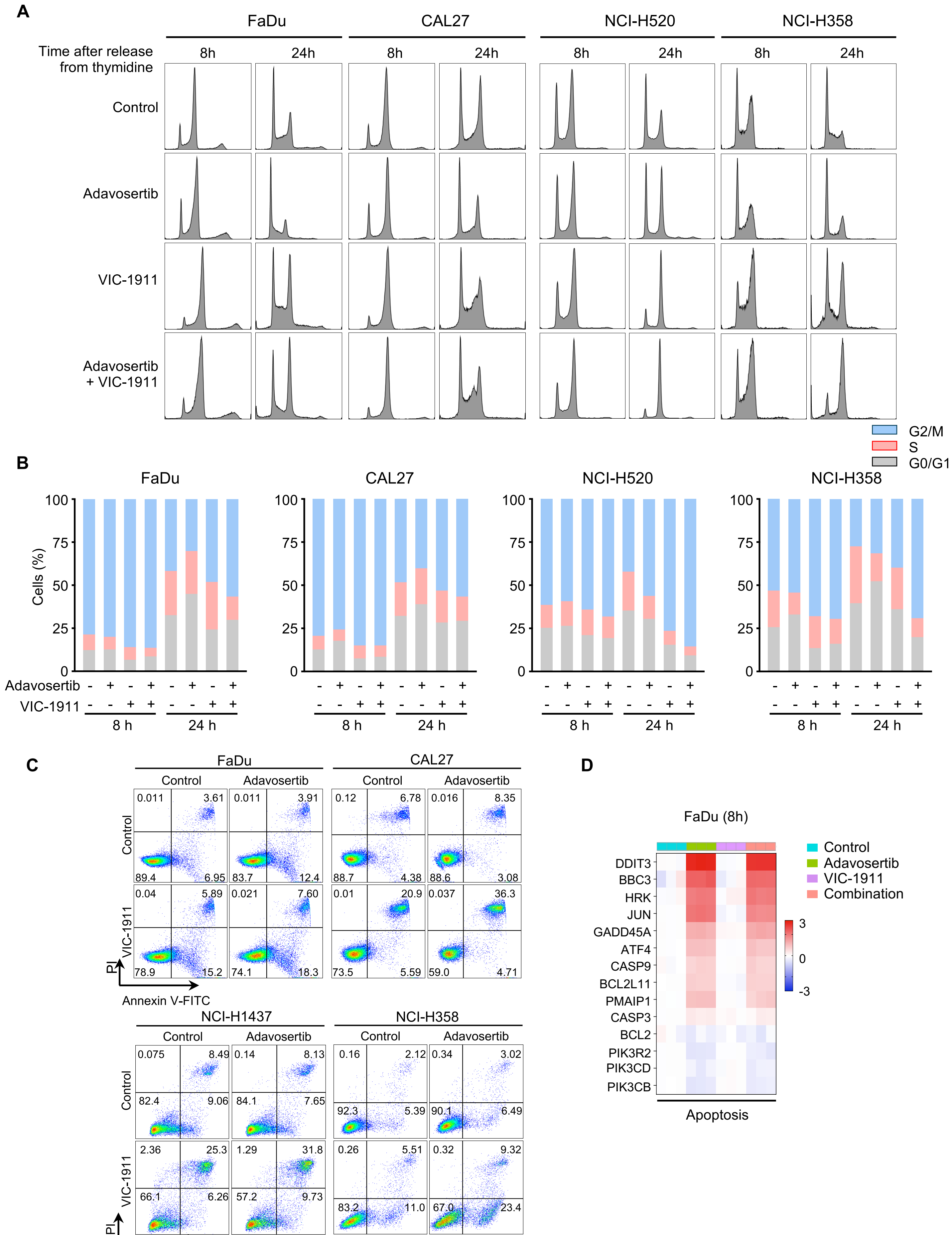


**Supplementary Figure S7**

**Supplementary Figure S7. Combination VIC-1911/adavosertib treatment increases cell proportion in G2/M through mitotic arrest and apoptosis. A** and **B**, Cell cycle distribution upon combined VIC-1911/adavosertib treatment in HNSCC and lung cancer cells. **A**, Cells were synchronized with double thymidine block, released and treated with control, 250 nmol/L adavosertib (500 nmol/L for CAL27), 125 nmol/L VIC-1911 (250 nmol/L for CAL27), or combination for 8h and 24h. Cell cycle distribution was assessed by flow cytometry following PI-staining. Representative images of cell cycle distribution in FaDu, CAL27, NCI-H520 and NCI-H358 cells. **B**, Quantification of cell population in the indicated cell cycle phases upon single or combination treatment. Data charts are representative of three independent experiments with consistent results. **C**, Representative images of flow cytometry to determine apoptotic cell death upon combinational treatment with VIC-1911 and adavosertib. HNSCC FaDu and CAL27, and lung cancer NCI-H1437 and NCI-H358 cells were treated with control, 250 nmol/L adavosertib, 125 nmol/L VIC-1911, or combination for 48h. Treated cells were further stained with FITC-conjugated Annexin V/PI and followed by flow cytometry to determine apoptotic cell death. Data images are representative of three independent experiments with consistent results. **D,** Heatmap showing increase of apoptotic genes in FaDu cells treated with the indicated drug for 8h.
