## Supplementary Figure S8 for "AURKA inhibition amplifies DNA replication stress to foster WEE1 kinase dependency and synergistic antitumor effects with WEE1 inhibition in cancers"

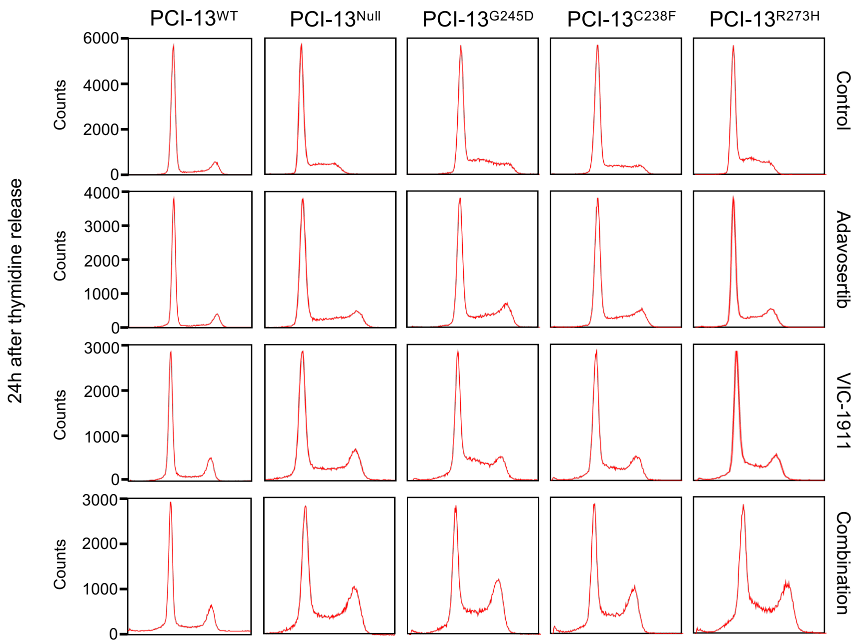


**Supplementary Figure S8. Combination AURKA/WEE1 exhibits higher S and G2/M cell population in *TP53*-mutated PCI-13 cells. Representative images of cell cycle distribution of the double-thymidine-**synchronized PCI-13 TP53 isogeneic cells after 24h exposure to combinational treatment VIC-1911/adavosertib for Fig. 5F.
