## Supplementary Figure S9 for "AURKA inhibition amplifies DNA replication stress to foster WEE1 kinase dependency and synergistic antitumor effects with WEE1 inhibition in cancers"

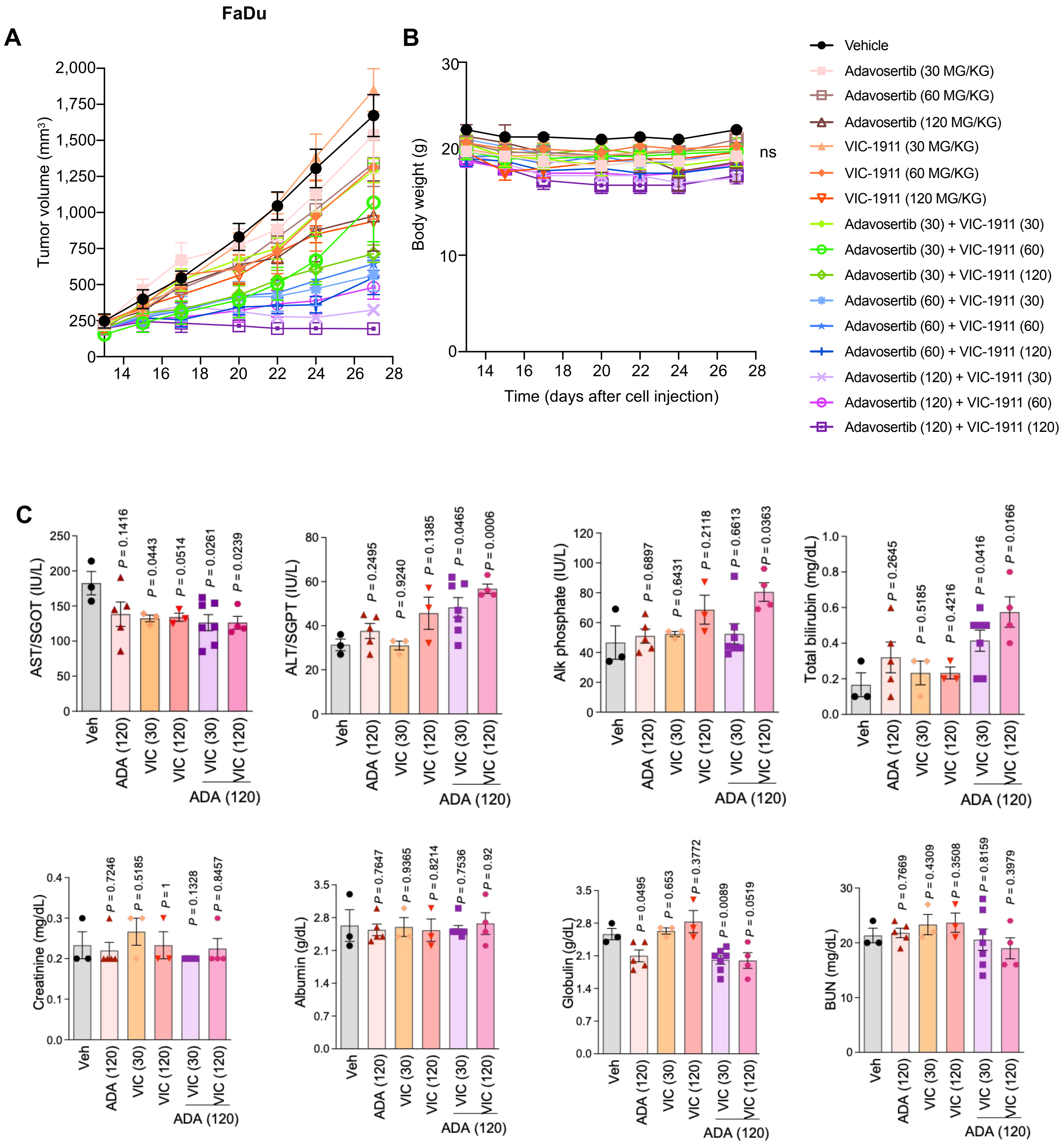


**Supplementary Figure S9**

**Supplementary Figure S9. Preclinical dose testing to optimize synergistic combination doses and toxicity profiling in HNSCC *in vivo* model. A** and **B,** FaDu CDX animals were daily treated with vehicle, adavosertib (30, 60, 120 mg $kg^{-1}$), VIC-1911 (30, 60, 120 mg kg-1) or 3 × 3 combinations through oral gavage for 14 days (6 days on-1 day off). Tumor volume (**A**) and body weight (**B**) were monitored every other day. Data are shown as mean ± SEM (n = 3-7 per group). **C**, Blood toxicity profile of VIC-1911/adavosertib treatment. Whole blood was collected at the endpoint of treatment duration in FaDu CDX animals and assessed by blood chemistry to determine toxicity profiles. Combinations VIC-1911 (30 mg $kg^{-1}$) + adavosertib (120 mg $kg^{-1}$) or VIC-1911 (120 mg $kg^{-1}$) + adavosertib (120 mg $kg^{-1}$) were selected for the toxicity charts including vehicle and single drugs. Data are shown as mean ± SEM and statistical significance (*P* value) are indicated.
