## Supplementary Figure S10 for "AURKA inhibition amplifies DNA replication stress to foster WEE1 kinase dependency and synergistic antitumor effects with WEE1 inhibition in cancers"

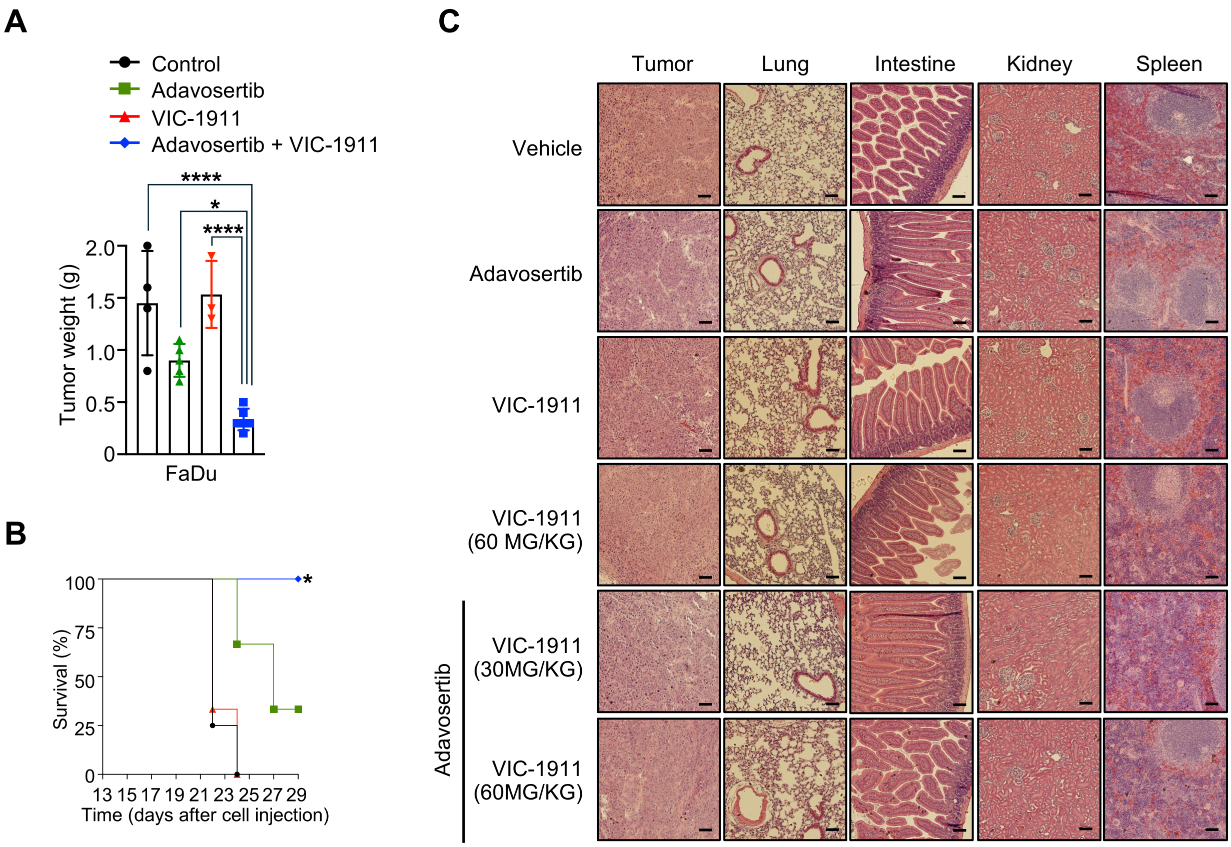


**Supplementary Figure S10. Combination VIC-1911/adavosertib treatment shows synergistic suppressive efficacy on tumors with no histopathological abnormalities in other organs. A** and **B,** Same study as presented in Fig. 6A with measurement of tumor weight (**A**) and survival curve of animals (**B**) in FaDu xenografted animals orally treated with control, 120 mg $kg^{-1}$ adavosertib, 30 mg $kg^{-1}$ VIC-1911, or in combination for 14 days. Data are shown as mean ± SEM. Statistic determination was applied a one-way ANOVA test. * $P < 0.05$, **** $P < 0.0001$. **C**, Representative images of histopathological evaluations in tumors and other organs including lung, intestine, kidney and spleen. Scale bar = 100 μm.
