## Supplementary Figure S11 for "AURKA inhibition amplifies DNA replication stress to foster WEE1 kinase dependency and synergistic antitumor effects with WEE1 inhibition in cancers"

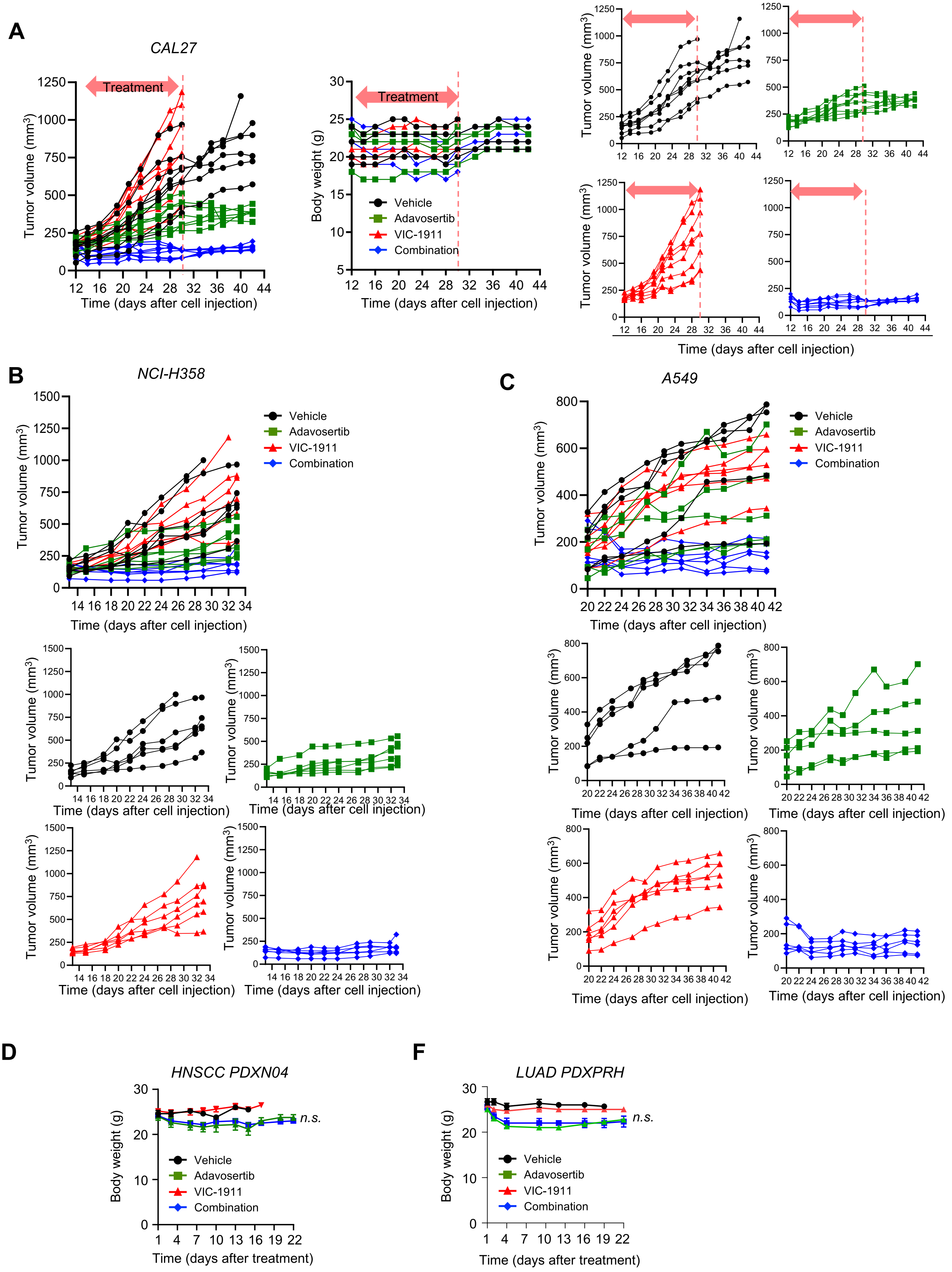


**Supplementary Figure S11**

**Supplementary Figure S11. Combined AURKA/WEE1 inhibition drives tumor regression in preclinical CDX and PDX models of HNSCC and lung cancer. A-C,** Same data as presented in Fig. 6B (**A**), Fig. 6C (**B**) and Fig. 6D (**C**) with tumor growth values of individual tumors in CAL27 (**A**), NCI-H358 (**B**) and A549 (**C**), respectively. Spider plots are indicated tumor growth of each tumor in the CDX models. **A**, CAL27 xenografted animals were orally treated with control, adavosertib, VIC-1911 or in combination for 18 days as treatment duration and further monitored tumor growth without treatment as indicated spider plots and body weight change. **B** and **C**, NCI-358 and A549 xenografted animals orally were treated with the indicated drug as single or combination for 21 days and tumor growth was presented as spider plots. **D** and **E**, Change in body weight in tumor-bearing PDX models including HNSCC PDXN04 (**D**) and LUAD PDXPRH (**E**) presented studies in Fig. 6F-6H and Fig. 6I-6K, respectively. PDXN04 and PDXPRH PDX animals were orally treated with control, 120 mg $kg^{-1}$ adavosertib, 30 mg $kg^{-1}$ VIC-1911, or combination and body weight change was monitored every other day for 21 days (6 days on-1 day off). *n.s.*; non-significance.
