## Supplementary Figure S12 for "AURKA inhibition amplifies DNA replication stress to foster WEE1 kinase dependency and synergistic antitumor effects with WEE1 inhibition in cancers"

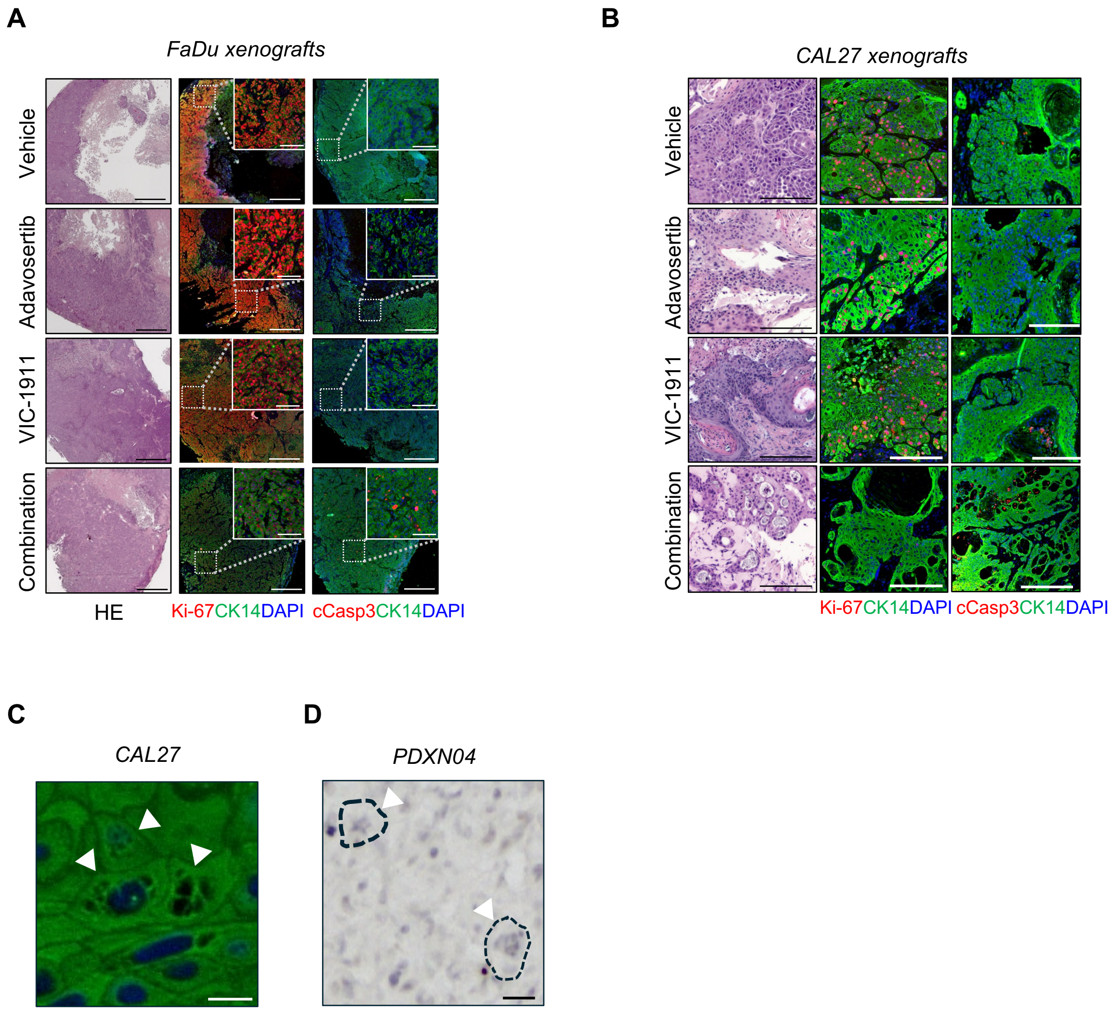


**Supplementary Figure S12. Combination VIC-1911/adavosertib treatment suppresses tumor cell proliferation and enhances apoptotic cell death in HNSCC CDX models. A** and **B,** Representative H&E and immunofluorescence stainings for CK14 (green) and either Ki-67 (red) or cleaved caspase 3 (red) in tumors in FaDu (**A**) and CAL27 (**B**) CDX animals treated with control, 120 mg $kg^{-1}$ adavosertib, 30 mg $kg^{-1}$ VIC-1911, or combination. Tumor tissues were harvested at 2h after treatment at the endpoint. Scale bar = 1 mm or 100 μm (inset). **C** and **D**, Representative images of micronuclei or multipolar spindles as arrow marked in CAL27 CDX (**C**) and PDXN04 PDX (**D**) tumor tissues treated with VIC-1911/adavosertib treatment. Dashed lines indicate cell margin. Scale bar = 10 μm.
