## Supplementary Table S1 for "AURKA inhibition amplifies DNA replication stress to foster WEE1 kinase dependency and synergistic antitumor effects with WEE1 inhibition in cancers"

### Supplementary Table 1. Clinical trials with AURKA inhibition

| Trial IDs | Drug(s) Used | Cancer Type | Reference | Status |
|---|---|---|---|---|
| ClinicalTrials.gov ID NCT05948475 | Tinengotinib | Fibroblast Growth Factor Receptor (FGFR)-altered, Chemotherapy- and FGFR Inhibitor-Refractory/Relapsed Cholangiocarcinoma | https://www.cancer.gov/research/participate/clinical-trials-search/v?id=NCI-2023-10661&r=1 | Yes |
| ClinicalTrials.gov ID NCT06457919 | Tinengotinib w/ enzalutamide or abiraterone acetate | Castration resistant Prostate Cancer | https://www.cancer.gov/research/participate/clinical-trials-search/v?id=NCI-2024-05117&r=1 | Yes |
| ClinicalTrials.gov ID NCT06095505 | Alisertib | Small Cell Lung Cancer (SCLC) following progression on or after treatment with one platinum-based chemotherapy and anti-PD-L1 immunotherapy agent | https://www.cancer.gov/research/participate/clinical-trials-search/v?id=NCI-2023-10166&r=1 | Yes |
| ClinicalTrials.gov ID NCT04085315 | Alisertib w/ Osimertinib | EGFR-mutated stage IV Lung Cancer | https://www.cancer.gov/research/participate/clinical-trials-search/v?id=NCI-2019-05913&r=1 | Yes |
| ClinicalTrials.gov ID NCT05490472 | JAB-2485 | Advanced solid tumors such as ER+ Breast Cancer, Triple Negative Breast Cancer (TNBC), AT-rich interaction domain 1A (ARID1A) Mutant Solid Tumors and Small Cell Lung Cancer (SCLC). | https://www.cancer.gov/research/participate/clinical-trials-search/v?id=NCI-2022-10634&r=1 | Yes |
| ClinicalTrials.gov ID NCT06369285 | Alisertib w/ Endocrine therapy | Pathology-confirmed HR-positive/HER2-negative Metastatic Breast Cancer (MBC) following progression on or after at least two prior lines of endocrine therapy | https://www.cancer.gov/research/participate/clinical-trials-search/v?id=NCI-2024-06850&r=1 | In Review |
| ClinicalTrials.gov ID NCT05374538 | VIC-1911 w/ Sotorasib | Locally advanced or metastatic KRAS G12C-mutant Non-small cell lung cancer (NSCLC) | https://www.cancer.gov/research/participate/clinical-trials-search/v?id=NCI-2022-05882 | Completed |
| ClinicalTrials.gov ID NCT05489731 | VIC-1911 w/ Osimertinib | Advanced Non-small Cell Lung Cancer (NSCLC) with EGRF- Mutation | https://ascopubs.org/doi/10.1200/JCO.2024.42.16_suppl.8077 | ? |
| ClinicalTrials.gov ID NCT01677559 | Alisertib w/ Nab-paclitaxel | Refractory high-grade Neuroendocrine tumours (NETs) | https://pubmed.ncbi.nlm.nih.gov/34256279/ | Completed |
| ClinicalTrials.gov ID NCT03092934 | AK-01 (LY3295668) | Locally advanced tumors Metastatic solid tumors Small Cell Lung Cancer (SCLC) Breast Cancer | https://pubmed.ncbi.nlm.nih.gov/33479856/ | Completed |
| ClinicalTrials.gov ID NCT01567709 | Alisertib w/ Vorinostat | Relapsed or Recurrent Hodgkin Lymphoma, B-Cell Non-Hodgkin Lymphoma, or Peripheral T-Cell Lymphoma | https://pubmed.ncbi.nlm.nih.gov/31617432/ | Completed |
