## Supplementary Table S2 for "AURKA inhibition amplifies DNA replication stress to foster WEE1 kinase dependency and synergistic antitumor effects with WEE1 inhibition in cancers"

### Supplementary Table 2. Clinical trials with WEE1 inhibition

| Trial IDs | Drug(s) Used | Cancer Type | Reference | Active? |
|---|---|---|---|---|
| ClinicalTrials.gov ID NCT04768868 | IMP7068 | Advanced Solid Tumors | https://www.cancer.gov/research/participate/clinical-trials-search/v?id=NCI-2021-13607&loc=0&q=WEE1&rl=1 | Yes |
| ClinicalTrials.gov ID NCT06457919 | Azenosertib (Zn-c3) w/ Gemcitabine | Advanced Pancreatic Cancer | https://www.cancer.gov/research/participate/clinical-trials-search/v?id=NCI-2023-09941&loc=0&q=WEE1&rl=1 | Yes |
| ClinicalTrials.gov ID NCT04516447 | Azenosertib (Zn-c3) | Ovarian Cancer | https://www.cancer.gov/research/participate/clinical-trials-search/v?id=NCI-2020-07262&loc=0&q=Zn-c3&rl=1 | Yes |
| ClinicalTrials.gov ID NCT05743036 | Azenosertib (Zn-c3) w/ encorafenib and cetuximab | Metastatic BRAF V600E mutant Colorectal Cancer | https://www.cancer.gov/research/participate/clinical-trials-search/v?id=NCI-2023-06212&loc=0&q=Zn-c3&rl=1 | Yes |
| ClinicalTrials.gov ID NCT06463340 | SGR-3515 | Advanced Solid Tumors | https://www.cancer.gov/research/participate/clinical-trials-search/v?id=NCI-2024-07565&loc=0&q=WEE1&rl=1 | Yes |
| ClinicalTrials.gov ID NCT06364410 | Azenosertib (Zn-c3) w/ Trastuzumab Deruxtecan (DS-8201a) | Locally advanced, metastatic, or unresectable HER2-positive gastric, gastroesophageal junction, or other solid tumors | https://www.cancer.gov/research/participate/clinical-trials-search/v?id=NCI-2024-02982&loc=0&q=WEE1&rl=1 | In review |
| ClinicalTrials.gov ID NCT06369155 | Azenosertib (Zn-c3) | Uterine Serous Cancer | https://www.cancer.gov/research/participate/clinical-trials-search/v?id=NCI-2024-04595&loc=0&q=WEE1&rl=1 | Temporarily closed to accrual |
| ClinicalTrials.gov ID NCT06351332 | Azenosertib (Zn-c3) w/ Carboplatin and Pembrolizumab | Metastatic Triple Negative Breast Cancer | https://www.cancer.gov/research/participate/clinical-trials-search/v?id=NCI-2024-03872&loc=0&q=WEE1&rl=1 | Temporarily closed to accrual |
| ClinicalTrials.gov ID NCT02659241 | Adavosertib | Primary advanced high grade serous ovarian, fallopian tube, or primary peritoneal cancer | https://www.cancer.gov/research/participate/clinical-trials-search/v?id=NCI-2016-00118&loc=0&q=WEE1&rl=1 | Temporarily closed to accrual |
| ClinicalTrials.gov ID NCT02482311 | Adavosertib | Ovarian Cancer (BRCA1/2 mutation [PARP-failures, Ovarian Cancer (BRCA wild-type) with more than three prior lines of treatment, Triple negative breast cancer (TNBC), and Small-cell lung cancer (SCLC) | https://pubmed.ncbi.nlm.nih.gov/37278879/ | Completed |
| ClinicalTrials.gov ID NCT01164995 | Adavosertib w/ Carboplatin | p53 mutated epithelial Ovarian Cancer | https://pubmed.ncbi.nlm.nih.gov/37236033/ | Completed |
